# F_1_F_O_-ATP synthase subunit DAPIT rewires mitochondrial metabolism in cancer cells to drive cellular hyperproliferation

**DOI:** 10.64898/2026.11.22.768406

**Authors:** Marie-Theres Borowski, Pia-Marie Ott, Martin Scholz, Michael Hippler, Karin B. Busch

## Abstract

Mitochondrial ATP synthase is the main source of cellular ATP. Its assembly into dimers and oligomers is modulated by several subunits. While subunits e and g promote dimerization, DAPIT (USMG5) stabilizes dimer-dimer interactions. Whether DAPIT (diabetes associated protein in insulin-sensitive tissues) also controls ATP synthase function remained unknown. Using high-resolution live-cell imaging, metabolic profiling, mass spectrometry, and molecular genetics methods, we show that DAPIT-dependent dimer stabilization delays enzyme turnover and optimizes ATP synthesis. This structural role extends beyond bioenergetics: by coupling mitochondrial energy conversion to metabolic restructuring, DAPIT enables the upregulation of glycolysis and efficient shunting of substrates toward oxidative phosphorylation. This allows cells to sustain energy metabolism without activating stress responses. Our findings establish DAPIT as a molecular link between ATP synthase architecture and metabolic adaptation, with implications for understanding mitochondrial remodeling in proliferating and differentiating cells.

**GRAPHICAL ABSTRACT:** 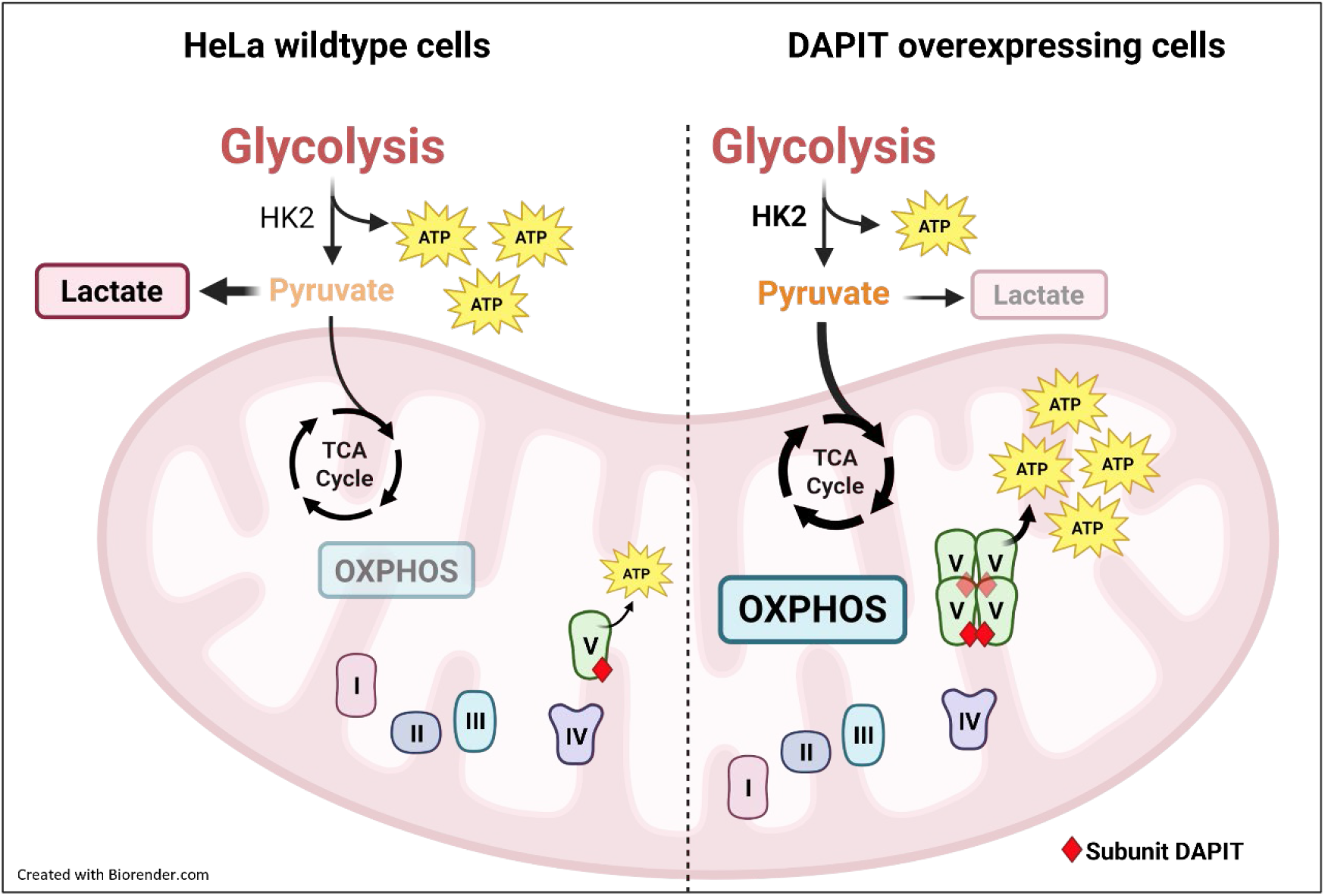

**IN BRIEF:** Borowski et al. demonstrate that the mitochondrial ATP synthase subunit DAPIT acts as a crucial metabolic switch. By stabilizing the ATP synthase and coupling enhanced hexokinase 2 (HK2) activity to mitochondrial OXPHOS, DAPIT rewires cellular metabolism. This resulting bioenergetic surplus accelerates cell cycle progression and drives cellular hyperproliferation.

**HIGHLIGHTS:**

- DAPIT stabilizes mitochondrial ATP synthase to optimize OXPHOS efficiency
- DAPIT rewires metabolism by coupling enhanced HK2 activity to OXPHOS
- This metabolic rewiring accelerates cell cycle progression and proliferation

## INTRODUCTION

Mitochondrial F_1_F_O_-ATP synthase is pivotal for mitochondrial and cellular physiology. Energy derived from oxidative metabolism is stored in a proton-motive force (PMF) across the inner mitochondrial membrane (IMM). The ATP synthase (complex V, CV, of the oxidative phosphorylation system) harnesses the PMF to generate ATP but also contributes to maintaining the PMF under unfavorable conditions via ATP hydrolysis (ATPase). This bidirectional activity places ATP synthase at the center of metabolic switches, allowing adaptation to rapid fluctuations in energy demand and nutrient availability. For example, during the transition from oxidative phosphorylation (OXPHOS) to glycolysis under hypoxia or in tumor cells, the reverse mode of ATP synthase is finely tuned to preserve mitochondrial membrane potential (ΔΨ_m_), thereby influencing cellular survival and metabolic reprogramming [1, 2]. Evidence is accumulating that the spatiotemporal organization of ATP synthase is critical for its function [3–5], but underlying mechanisms are still unresolved.

Mitochondrial F_1_F_O_-ATP synthase is organized in dimers that form rows along the rims of cristae. This supramolecular organization is critical for shaping and stabilizing cristae architecture [6, 7], acting together with OPA1 (Optic atrophy 1) and the MICOS (mitochondrial contact site and cristae organizing system) complex to define mitochondrial morphology and function [8, 9].

Dimerization of F_1_F_O_-ATP synthase is primarily promoted by subunits e and g, while subunits IF1 and DAPIT contribute to higher-order stabilization of dimer–dimer interfaces [10, 11]. DAPIT (diabetes-associated protein in insulin-sensitive tissues) is a small, 58-amino acid, 6.4 kDa single-pass transmembrane protein encoded by the *USMG5* gene [12, 13]. It is evolutionarily conserved from insects to vertebrates [12, 14] and is upregulated during skeletal muscle growth [13], suggesting an essential physiological role. Structurally, DAPIT is localized at the peripheral outermost edge of the F_O_ complex and is incorporated during the final steps of assembly [15, 16]. Recent structural studies of porcine ATP synthase tetramers and bovine ATP synthase dimers [17–19] place DAPIT at dimer–dimer contact sites, and loss of DAPIT destabilizes both CV and its dimeric forms [11, 16, 20]. Interestingly, mammalian DAPIT is annotated as an ortholog of subunit k in *Saccharomyces cerevisiae* (ATPMK) [21], yet sequence and structural similarity between these proteins is limited, leaving the molecular function of DAPIT incompletely understood. Emerging evidence, however, underscores its critical physiological role in tissues with high bioenergetic and metabolic demands. For instance, DAPIT expression is induced by glucose in pancreatic β-cells [22], and its dysregulation is a key feature of the mitochondrial dysfunction observed in Duchenne muscular dystrophy [23]. Furthermore, loss of DAPIT directly induces dilated cardiomyopathy in zebrafish models [24], and initial *in vitro* studies suggest that overexpressing DAPIT can modulate cellular glucose metabolism [25].

Despite its evolutionary conservation and emerging structural importance, the precise mechanistic role of DAPIT in ATP synthase dimer stabilization and mitochondrial physiology remains an open question. Here, we show how DAPIT, as a small peripheral subunit, stabilizes ATP synthase dimers, thereby delaying the degradation of ATP synthase and optimizing the efficiency of ATP synthesis. This process couples mitochondrial energy conversion to metabolic restructuring, leading to upregulation of glycolysis and efficient substrate shunting to enhance OXPHOS without inducing cellular stress.

## RESULTS

Structural modeling based on ovine ATP synthase monomer [26] visualized that the membrane-embedded DAPIT subunit is positioned at the interface of the rotor-stator interface of F_O_ domain (Figure 1A). However, the structure of porcine ATP synthase tetramer [17] suggests that individual DAPIT subunits are too distant from one another to facilitate direct self-interaction (Figure 1B). This structural arrangement was further supported by an *in silico* structure prediction of human DAPIT sequence using AlphaFold 3 (Figure S1A, [27]). To experimentally dissect the functions of DAPIT, we generated and validated HeLa DAPIT overexpression (OE) and DAPIT knockout (KO) cell lines. Successful modulation of DAPIT expression was confirmed via Western blot analysis of cell lysates, showing a significant increase in DAPIT protein level in DAPIT OE cells and a complete absence in DAPIT KO cells (Figure 1C). To verify the sub-cellular localization of overexpressed DAPIT, we performed Western blot analysis of isolated membrane fractions and purified mitochondria, demonstrating its successful targeting and enrichment within the mitochondria (Figure S1B, C). As the glyceraldehyde 3-phosphate dehydrogenase (GAPDH) protein levels across all cell lines were stable, it was used for normalization (Figure S1D). With these cell models we investigated the structural and functional effects of varying DAPIT levels on the mitochondrial OXPHOS system.

**Figure 1:**
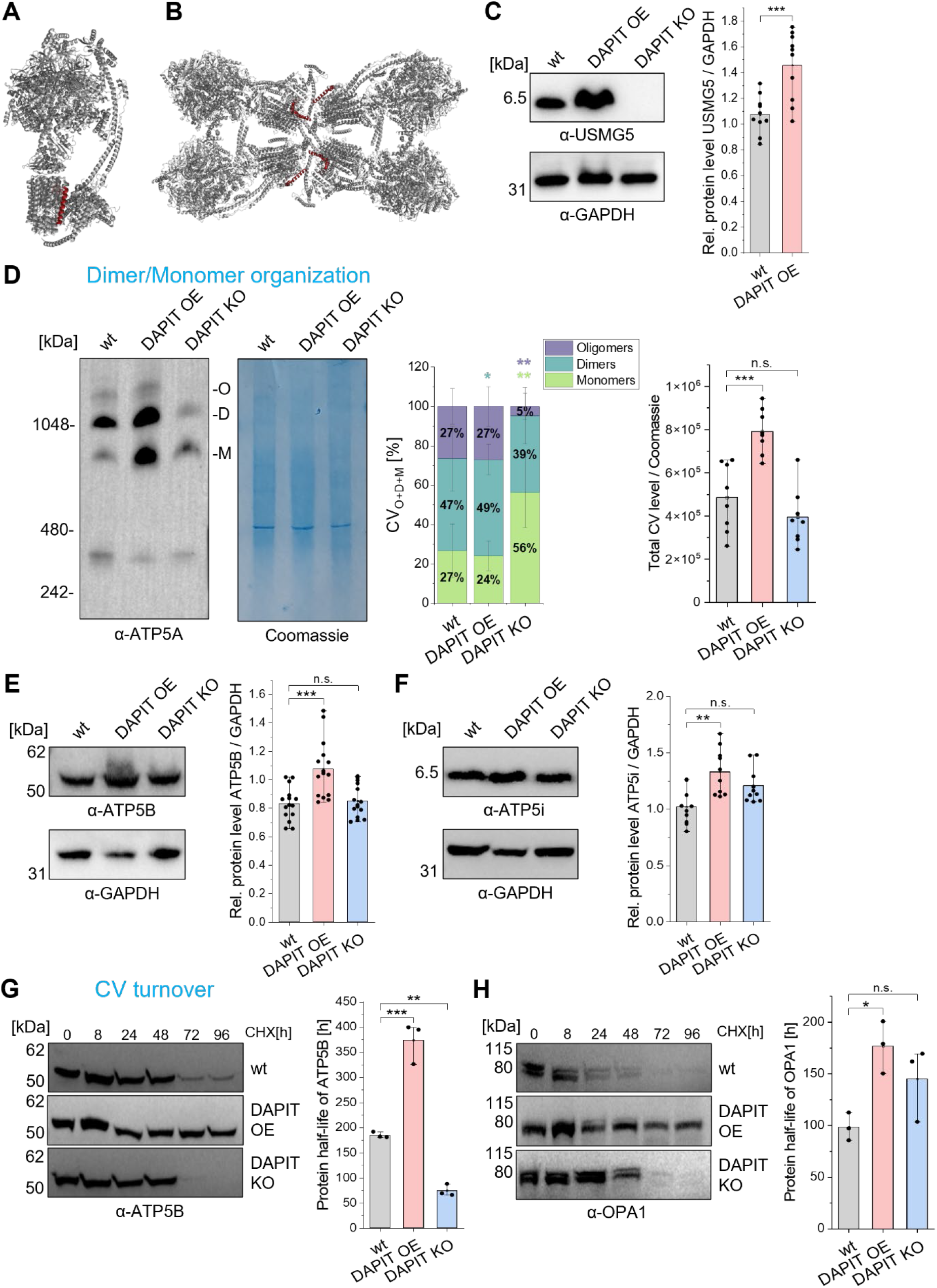
Elevated DAPIT levels promote ATP synthase oligomerization and stability. **(A)** Ovine ATP synthase monomer structure (PDB ID: 6TT7 [16]) with membrane-embedded DAPIT subunit highlighted in red. **(B)** Porcine ATP synthase tetramer structure (PDB ID: 6J5K [13]) with membrane-embedded DAPIT subunit highlighted in red. **(C)** Representative Western blot (probed with α-USMG5 and α-GAPDH) and quantification of relative protein (rel.) levels of USMG5 (also known as ATP synthase subunit DAPIT) in HeLa wildtype (wt), DAPIT overexpressing (OE), and DAPIT knockout (KO) cells normalized to GAPDH (biological experiments N=5, with each technical replicates n=2; one-way ANOVA with Scheffé’s *post-hoc* test). **(D)** Representative clear-native PAGE immunoblotted for subunit α (α-ATP5A), showing monomers (M), dimers (D), and oligomers (O) of ATP synthase from isolated mitochondria. Quantification of oligomerization state of ATP synthase (complex V, CV) reveals decreased oligomeric forms in DAPIT KO cells, and increased dimers and total CV levels in DAPIT OE cells normalized to Coomassie staining (N=4, n=2; one-way ANOVA with Scheffé’s *post-hoc* test; independent mitochondrial isolations). **(E, F)** Representative Western blots and quantification of relative protein levels of **(E)** ATP synthase subunit β (α-ATP5B) and **(F)** subunit e (α-ATP5i) normalized to GAPDH (α-ATP5B N=5, n=3; one-way ANOVA with Scheffé’s *post-hoc* test; α-ATP5i N=3, n=3; Kruskal-Wallis with Dunn’s *post-hoc* test). **(G, H)** Representative Western blots and quantification of relative protein levels of **(G)** ATP synthase subunit β (α-ATP5B) and **(H)** OPA1 after 0, 8, 24, 48, 72, and 96 h of cycloheximide (CHX) treatment. Equal amounts of total protein (25 µg protein) were loaded per lane. Protein half-lives were calculated via linear regression of relative band intensities (normalized to t_0_=100%) plotted against time (N=3, n=1; one-way ANOVA with Scheffé’s *post-hoc* test). Data are presented as mean ± SD of N=3-5 independent biological experiments with overlaid individual data points. The exact number of independent experiments (N) and each number of analyzed technical replicates, wells, or individual cells (n) are indicated in the respective brackets for each panel. Statistics: ***p≤0.001; **p≤0.01; *p≤0.05. (See also Figure S1). ANOVA, analysis of variance; DAPIT, diabetes-associated protein in insulin-sensitive tissue; GAPDH, glyceraldehyde-3-phosphate dehydrogenase; OPA1, optic atrophy 1; SD, standard deviation; USMG5, up-regulated during skeletal muscle growth protein 5.

### DAPIT modulates the oligomerization state and protein stability of human ATP synthase

Since DAPIT is localized in the membrane domain of human ATP synthase, we first analyzed whether its OE or absence directly impacts the oligomerization of ATP synthase. Clear-native (CN) PAGE followed by immunoblotting against subunit α (α-ATP5A) allowed the separation of ATP synthase monomers (M), dimers (D), and oligomers (O) (Figure 1D). Quantitative analysis revealed that DAPIT KO cells exhibited a significant loss of oligomeric forms, resulting in an increase of monomers. Conversely, DAPIT OE cells displayed a significant increase in dimeric ATP synthase forms and total ATP synthase levels. These changes occurred alongside alterations in protein levels of other key ATP synthase subunits, as the relative protein levels of subunit β (α-ATP5B) and subunit e (α-ATP5i) remained significantly increased in DAPIT OE cells (Figure 1E, F). To determine whether the structural stabilization of ATP synthase in the presence of abundant DAPIT impacts its degradation kinetics, we assessed protein turnover using a cycloheximide (CHX) chase assay over a time course of up to 96 hours. The protein half-life of core subunit β (α-ATP5B) was significantly prolonged in DAPIT OE cells compared to wildtype cells (Figure 1G). In addition, the protein half-life of the mitochondrial fusion OPA1 was also significantly increased in DAPIT OE cells, suggesting that the overexpression of DAPIT promotes mitochondrial structural stability (Figure 1H). In contrast, DAPIT KO cells showed a significant reduction in protein half-life of subunit β (α-ATP5B) (Figure 1G), indicating that the loss of DAPIT destabilizes the molecular structure, thereby accelerating the turnover of key subunits of the F_1_ domain. Taken together, these results show that while DAPIT is not necessarily required for the expression of individual core subunits [16] in HAP1 DAPIT KO cells, it nevertheless plays a crucial role in stabilizing the higher-order oligomerization states and in regulating the protein turnover of ATP synthase complex.

### Elevated DAPIT expression optimizes mitochondrial respiration and bioenergetic efficiency

Considering that DAPIT regulates the oligomerization and stability of ATP synthase, we next investigated whether these structural alterations led to functional changes in bioenergetic efficiency. We first evaluated the catalytic capacity of ATPase using an in-gel ATP hydrolysis activity assay following CN-PAGE separation (Figure 2A, [28]). Quantification of the resulting white lead phosphate precipitates, normalized to relative subunit β (α-ATP5B) levels, revealed that total ATP hydrolysis capacity was significantly decreased in DAPIT OE cells and remained unchanged in DAPIT KO cells. To determine whether the mitochondrial ATPase inhibitor protein 1 (IF1) could modulate this activity, we quantified its protein levels via Western blot (Figure 2B). The relative protein levels of IF1 (α-ATPIF1) were significantly decreased in both DAPIT OE and DAPIT KO cells compared to wildtype cells. To comprehensively map the cellular bioenergetic profile, we monitored oxygen consumption rates (OCR) and proton efflux rates (PER), calculated from extracellular acidification rates, using a Seahorse XF96 Flux Analyzer (Figure 2C and Figure S2A). By sequentially applying specific metabolic inhibitors – oligomycin (ATP synthase inhibitor), rotenone (complex I inhibitor), and antimycin A (complex III inhibitor) – we quantified the total ATP production rates derived from both glycolysis (glycoATP) and OXPHOS (mitoATP) (Figure 2C). DAPIT OE cells displayed an enhancement in mitoATP, glycoATP, and total ATP production rates (totalATP), whereas DAPIT KO cells exhibited no differences. To distinguish these bioenergetic effects from broader cellular signaling and to confirm a direct mitochondrial optimization, we next isolated intact mitochondria and kinetically traced ATP production rates using a luciferase assay (Figure 2D and Figure S2B). Succinate and rotenone were provided to fuel respiration exclusively via complex II, while adenylate kinase was blocked with Ap5A. To ensure quantification of specific mitochondrial ATP synthesis, the basal slope was corrected by subtracting the remaining background slope obtained in the presence of ATP synthase inhibitor oligomycin (Figure S2B). The final oligomycin-sensitive mitochondrial ATP production rates were significantly accelerated in mitochondria isolated from DAPIT OE cells, whereas DAPIT KO mitochondria showed no changes in ATP synthesis kinetics (Figure 2D). To investigate whether this increased ATP synthesis in DAPIT OE cells alters oxygen consumption and OXPHOS activity, we performed a Seahorse Mito Stress Test. DAPIT OE cells showed a significant increase in both basal and maximal respiration, as well as in non-mitochondrial respiration (Figure 2E and Figure S2C). In contrast, DAPIT depletion caused a reduction in OCR. To rule out that this altered metabolic state results from a change in mitochondrial uncoupling systems, we quantified the relative mRNA expression of uncoupling proteins 2 and 3 (*UCP2* and *UCP3*) via quantitative real-time PCR (Figure 2F). Expression levels of both *UCP2* and *UCP3* were significantly decreased in DAPIT OE cells, suggesting minimized proton loss to maximize the efficiency of ATP synthesis. Conversely, DAPIT KO cells exhibited an upregulation of *UCP2*, which likely results in respiratory inefficiency. Since a reduction in proton uncoupling directly influences the electrochemical gradient across the IMM, we next monitored ΔΨ_m_ in live cells using fluorescence microscopy (Figure 2G and Figure S2D). To account for variations in mitochondrial network size, the TMRE signal intensity was normalized to mitochondrial mass using the TMRE/MitoTracker Green (MTG) ratio. DAPIT OE cells exhibited a significantly increased ΔΨ_m_ compared to wildtype cells, which was sensitive to the addition of ATP synthase inhibitors oligomycin or venturicidin A, similar to the responses observed in wildtype cells (Figure 2G). In contrast, DAPIT KO cells exhibited an unchanged ΔΨ_m_, which continued to respond to venturicidin A but remained insensitive to oligomycin treatment. Since oligomycin reduced oxygen consumption in DAPIT KO cells (Figure 2E), this insensitivity likely is not due to a failure of inhibitor binding. Rather, this could indicate altered proton flux dynamics. The significant upregulation of *UCP2* expression in DAPIT KO cells (Figure 2F) likely functions as an active bypass. When ATP synthase is blocked by oligomycin, protons can flow through this *UCP2* leak, thereby preventing a shift in ΔΨ_m_. In addition, ΔΨ_m_ was fully abolished upon addition of the uncoupler CCCP in all cell lines (Figure S2D). Overall, these bioenergetic analyses demonstrate that DAPIT is a key factor in mitochondrial efficiency. Its abundance directly affects OXPHOS activity, respiratory coupling, and ATP synthesis.

**Figure 2:**
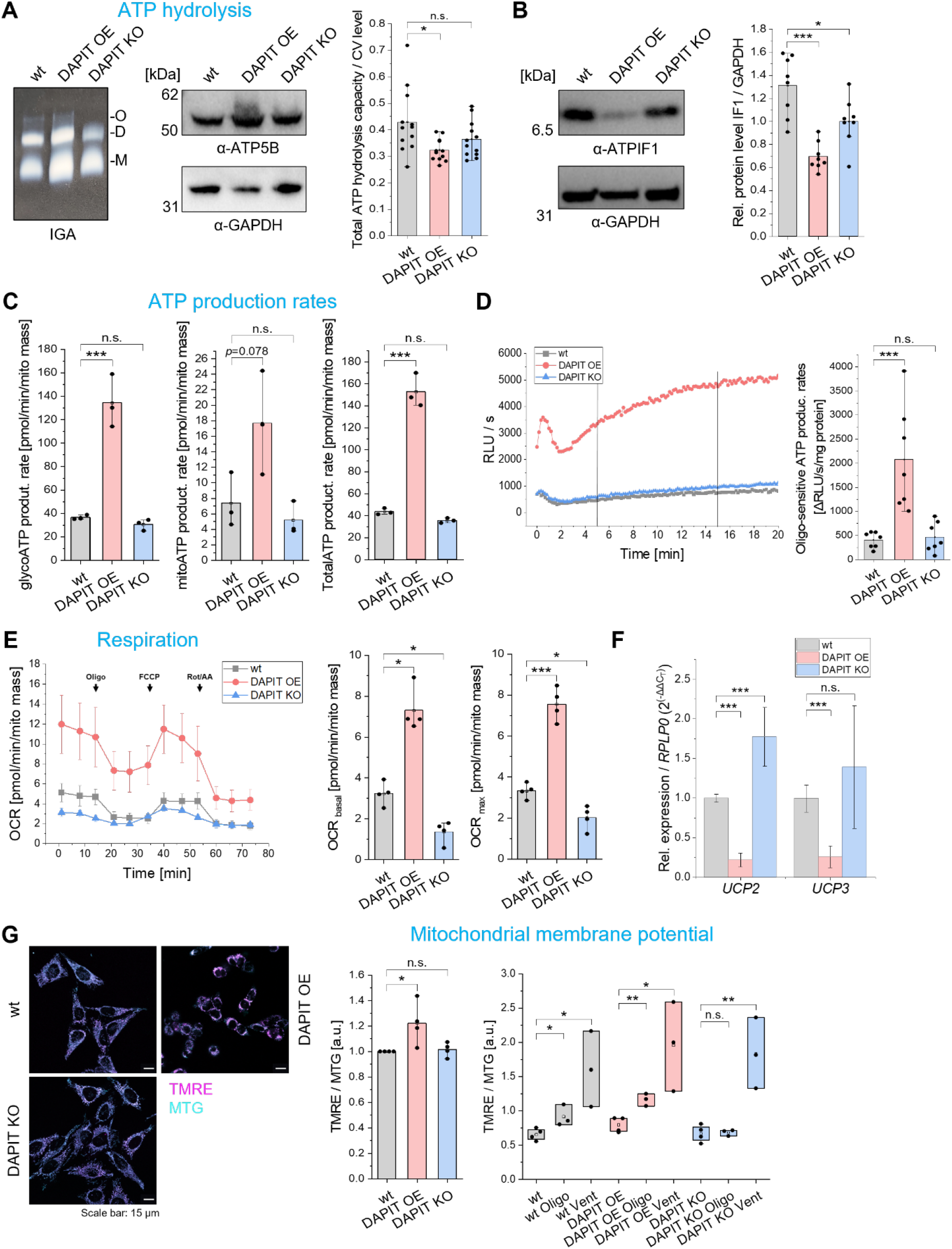
DAPIT stimulates mitochondrial respiration and oxidative coupling to drive ATP synthesis. **(A)** Representative in-gel activity (IGA) assay for determining ATP hydrolysis function, in which ATP synthase complexes were separated using clear-native PAGE. ATP hydrolysis correlates with development of white lead phosphate precipitates. Total ATP hydrolysis capacity was normalized to ATP synthase (CV) levels; representative Western blot showing ATP synthase subunit β (α-ATP5B) protein levels (N=4, n=3 one-way ANOVA with Scheffé’s *post-hoc* test). **(B)** Representative Western blot and quantification of relative protein (rel.) levels of IF1 (α-ATPIF1) normalized to GAPDH (N=4, n=2; one-way ANOVA with Scheffé’s *post-hoc* test). **(C)** ATP production (product.) rates were determined using oxygen consumption rates and extracellular acidification rates measured with an automatic Seahorse XF96 flux analyzer. By subsequently adding ATP synthase inhibitor oligomycin (1 μM), complex I inhibitor rotenone (1 μM), and complex III inhibitor antimycin A (1 μM), ATP production rates from glycolysis (glycoATP), oxidative phosphorylation (mitoATP), and total ATP (totalATP) were quantified and normalized to mitochondrial mass (mito mass) (N=3, n=20 wells; one-way ANOVA with Scheffé’s *post-hoc* test). **(D)** Representative kinetic profile of ATP production rates from freshly isolated mitochondria of HeLa wildtype (wt), DAPIT overexpressing (OE), and DAPIT knockout (KO) cells, measured as RLU/s using a luciferase assay. Equal amounts of total protein (50 µg mitochondria) were used. ATP production was fueled via complex II by addition of succinate (5 mM) in presence of rotenone (5 mM) to inhibit complex I. ADP (4 mM) was added as a further substrate and adenylate kinase was inhibited with Ap5A (20 µM). Quantification of ATP production rates was calculated from the slope of linear phase between minutes 5-15 (black lines). To determine specific mitochondrial ATP production, the slope in presence of oligomycin (10 µM) was subtracted from the basal slope. Final oligomycin (Oligo)-sensitive ATP production rates were normalized to mitochondrial protein input (N=3, n=2 wells; one-way ANOVA with Scheffé’s *post-hoc* test; independent mitochondrial isolations). **(E)** Functional test of mitochondrial respiration with subsequent addition of ATP synthase inhibitor oligomycin (Oligo, 1 μM), uncoupler FCCP (1 μM), complex I inhibitor rotenone (Rot, 1 μM), and complex III inhibitor antimycin A (AA, 1 μM), as indicated by arrows. Oxygen consumption rates (OCR) of HeLa wt, DAPIT OE, and DAPIT KO cells were determined with an automatic Seahorse XF96 flux analyzer and normalized to mitochondrial mass (mito mass). Basal and maximal (max) respiration was determined (N=4, n=20 wells; basal Kruskal-Wallis with Dunn’s *post-hoc* test; max one-way ANOVA with Scheffé’s *post-hoc* test). **(F)** Quantification of relative mRNA expression levels of uncoupling protein 2 (*UCP2*) and *UCP3*. Expression levels were determined by quantitative real-time PCR and normalized to housekeeping gene *RPLP0* using the 2^-ΔΔCt^ method with control group (wt) set to 1 (fold change) (*UCP2* N=4, n=3; Kruskal-Wallis with Dunn’s *post-hoc* test; *UCP3* N=5, n=3; one-way ANOVA with Scheffé’s *post-hoc* test). **(G)** Representative overlaid live-cell fluorescence microscopy images showing mitochondrial membrane potential (ΔΨ_m_) stained with TMRE (magenta) and mitochondrial mass stained with MitoTracker Green (MTG, cyan). Mitochondrial membrane potential was calculated as the TMRE/MTG ratio with control group (wt) set to 1 (fold change). As control, ATP synthase inhibitors oligomycin (Oligo, 5 µM) and venturicidin A (Vent, 5 µM) were added after staining (N=4, n=35 single cells; oligo N=3, n=21 single cells; vent N=3, n=23 single cells; one-way ANOVA with Scheffé’s *post-hoc* test). Data are presented as mean ± SD of N=3-4 independent biological experiments. The exact number of independent experiments (N) and each number of analyzed technical replicates, wells, or individual cells (n) are indicated in the respective brackets for each panel. Individual data points are overlaid in (A), (C), and (D). Statistics: ***p≤0.001; **p≤0.01; *p≤0.05. (See also Figure S2). ANOVA, analysis of variance; Ap5A, diadenosine pentaphosphate; DAPIT, diabetes-associated protein in insulin-sensitive tissue; FCCP, carbonyl cyanide 4-(trifluoromethoxy)phenylhydrazone; IF1, ATPase inhibitory factor 1; RLU, relative light units; RPLP0, ribosomal protein lateral stalk subunit P0; SD, standard deviation; TMRE, tetramethylrhodamine ethyl ester.

### DAPIT abundance regulates mitochondrial network architecture and cellular morphology

After we determined that DAPIT is necessary for the oligomerization of ATP synthase and for efficient OXPHOS activity, we next investigated whether these modulations translate into alterations in the mitochondrial network and in overall cell morphology.

Live-cell fluorescence microscopy using MTG to stain mitochondria and subsequent quantitative analysis revealed that the relative mitochondrial area normalized to cell size was significantly decreased in both DAPIT OE and DAPIT KO cells (Figure 3A). In DAPIT OE cells, this reduction in mitochondrial mass was confirmed by a significant decrease in relative protein levels of VDAC (Figure S3A). Consistent with the reduced mitochondrial mass, this structural remodeling in DAPIT OE cells was accompanied by a transcriptional downregulation of the mitochondrial biogenesis regulator peroxisome proliferator-activated receptor gamma coactivator 1-alpha (*PGC1α*) (Figure 3B). In contrast, DAPIT KO cells showed a significant increase in relative expression levels of PGC1α, suggesting a compensatory biogenesis response to the loss of functional mitochondria. These changes in mitochondrial mass in DAPIT OE cells were accompanied by significant changes in cell or nuclear size. Quantification of cell boundaries and nuclear areas showed that DAPIT OE cells were significantly smaller and exhibited smaller nuclei compared to wildtype cells (Figure 3C, and S3B). In contrast, DAPIT KO cells displayed no alterations in cell and nuclear size. To exclude the possibility that these alterations in cell and nuclei size in DAPIT OE cells were consequences of activation of cellular stress pathways, we analyzed the expression of activating transcription factor 4 (ATF4), a marker for integrated stress response. Western blot analysis revealed that relative ATF4 protein levels in DAPIT OE cells remained at wildtype levels (Figure S3C), confirming that overexpression of DAPIT does not trigger a pathological stress response despite the induced morphological changes.

**Figure 3:**
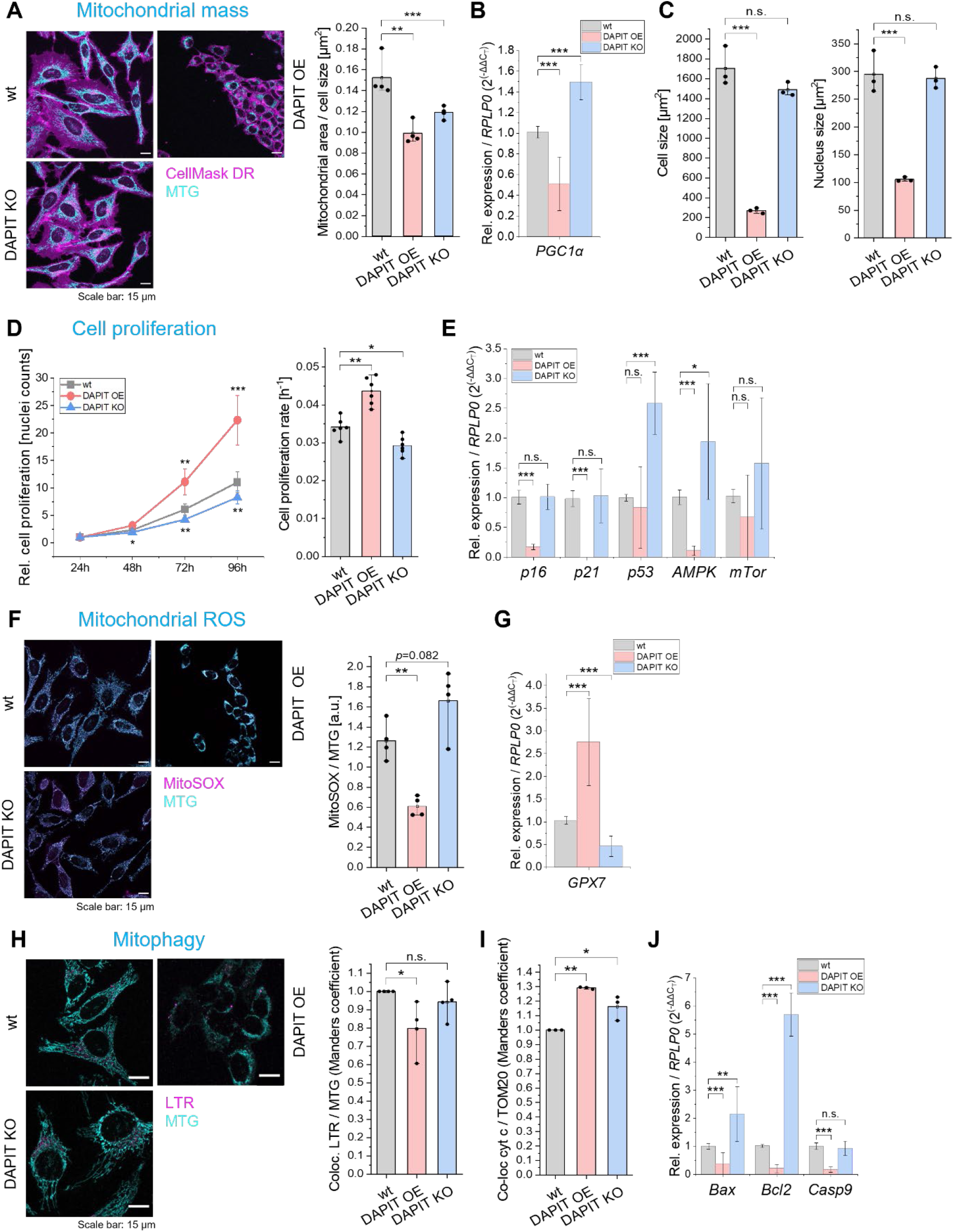
DAPIT regulates mitochondrial network integrity to sustain cellular proliferation and prevent oxidative stress. **(A)** Representative overlaid images from live-cell fluorescence microscopy showing plasma membrane staining with CellMask Deep-Red (DR, magenta) and MitoTracker Green (MTG, cyan) to determine mitochondrial mass. Mitochondrial area was determined using the mitochondrial network analysis plugin (MiNA) [29] for Fiji and the ratio of mitochondrial area to cell size was calculated (N=4, n=32 single cells; Kruskal-Wallis with Dunn’s *post-hoc* test). **(B)** Quantification of relative (rel.) mRNA expression levels of *PGC1α*. Expression levels were determined by quantitative real-time PCR and normalized to housekeeping gene *RPLP0* using the 2^-ΔΔCt^ method with control group (wt) set to 1 (fold change) (N=4, n=3; Kruskal-Wallis with Dunn’s *post-hoc* test). **(C)** Quantification of cell size and nucleus size by staining plasma membrane with CellMask DR and nuclei with Hoechst 33342 (cell size N=3, n=25; nucleus size N=3, n=31 single cells; one-way ANOVA with Scheffé’s *post-hoc* test). **(D)** Growth curves of HeLa wildtype (wt), DAPIT overexpressing (OE), and DAPIT knockout (KO) cells over a period of 96 h. Cell numbers were determined every 24 h by counting Hoechst 33342-stained nuclei using a fluorescence microplate reader. To account for initial seeding differences, cell numbers were normalized to 24 h timepoint. Cell proliferation rates were calculated from the slope of exponential growth phase (N=3, n=2; one-way ANOVA with Scheffé’s *post-hoc* test). **(E)** Quantification of relative mRNA expression levels of *p16*, *p21*, *p53*, *AMPK*, and *mTOR*. Expression levels were determined by quantitative real-time PCR and normalized to housekeeping gene *RPLP0* using the 2^-ΔΔCt^ method with control group (wt) set to 1 (fold change) (*p16* and *p21* N=4, n=3; *p53* and *AMPK* N=5, n=3; *mTOR* N=8, n=24; Kruskal-Wallis with Dunn’s *post-hoc* test). **(F)** Representative overlaid images from live-cell fluorescence microscopy showing staining of mitochondrial reactive oxygen species with MitoSOX (magenta) and mitochondrial staining with MitoTracker Green (MTG, cyan). Mitochondrial reactive oxygen species levels were normalized to mitochondrial mass (N=4, n=30 single cells; one-way ANOVA with Scheffé’s *post-hoc* test). **(G)** Quantification of relative mRNA expression levels of *GPX7*. Expression levels were determined by quantitative real-time PCR and normalized to housekeeping gene *RPLP0* using the 2^-ΔΔCt^ method with control group (wt) set to 1 (fold change) (N=5, n=3; one-way ANOVA with Scheffé’s *post-hoc* test). **(H)** Representative overlaid images from live-cell fluorescence microscopy showing staining of lysosomes with LysoTracker Red (LTR, magenta) and mitochondria with MitoTracker Green (MTG, cyan). Mitophagy was determined by quantifying colocalization (coloc) of lysosomes with mitochondria (Manders’ coefficient) using the Coloc 2 plugin for Fiji, with control group (wt) set to 1 (fold change) (N=4, n=40 single cells; Kruskal-Wallis with Dunn’s *post-hoc* test). **(I)** Determination of colocalization (coloc) of cytochrome c (cyt c) with TOM20 as marker for mitochondria, using the Coloc 2 plugin for Fiji after immunofluorescence staining. Loss of cytochrome c/TOM20 colocalization serves as an indicator of mitochondrial outer membrane permeabilization and apoptotic cytochrome c release. Control group (wt) was set to 1 (fold change) (N=3, n=40 single cells; one-way ANOVA with Scheffé’s *post-hoc* test). **(J)** Quantification of relative mRNA expression levels of *Bax*, *Bcl2*, and *Casp9*. Expression levels were determined by quantitative real-time PCR and normalized to housekeeping gene *RPLP0* using the 2^-ΔΔCt^ method with control group (wt) set to 1 (fold change) (*Bax* N=4, n=3; *Bcl2* and *Casp9* N=3, n=3; Kruskal-Wallis with Dunn’s *post-hoc* test). Data are presented as mean ± SD of N=3-5 independent biological experiments. The exact number of independent experiments (N) and each number of analyzed technical replicates, wells, or individual cells (n) are indicated in the respective brackets for each panel. Individual data points are overlaid in (D). Statistics: ***p≤0.001; **p≤0.01; *p≤0.05. (See also Figure S3). AMPK, AMP-activated protein kinase; ANOVA, analysis of variance; Bax, Bcl2-associated X protein; Bcl2, B-cell lymphoma 2; Casp9, caspase 9; DAPIT, diabetes-associated protein in insulin-sensitive tissue; GPX7, glutathione peroxidase 7; mTOR, mechanistic target of rapamycin; PGC1α, peroxisome proliferator-activated receptor gamma coactivator 1-α; RPLP0, ribosomal protein lateral stalk subunit P0; SD, standard deviation; TOM, translocase of the outer membrane.

### DAPIT abundance determines cell proliferation rates

This data in combination with the bioenergetic analyses (Figure 2) suggest that DAPIT OE yields fewer but more efficient mitochondria, whereas its absence triggers mitochondrial and respiratory deficit. Since cell division is dependent on cellular energy state, we next examined how these different phenotypes affect overall cellular fitness. First, we monitored cell proliferation over 96 hours. DAPIT OE cells showed a significantly increased cell proliferation rate compared to wildtype cells. In contrast, DAPIT KO cells exhibited a decreased cell proliferation rate (Figure 3D). A similar cell proliferation phenotype was also observed under OXPHOS conditions (galactose medium) (Figure S4A), which rules out that this effect is driven by glycolytic conditions (high-glucose medium). To understand the molecular pathways underlying these divergent growth phenotypes, we analyzed the transcriptional expression of key cell cycle inhibitors and metabolic sensors (Figure 3E). In DAPIT OE cells, the enhanced cell proliferation rate was accompanied by a significant transcriptional downregulation of cell cycle arrest genes *p16* and *p21*, while *p53* expression levels remained unchanged. At the same time, the expression of the energy sensor AMP-activated protein kinase (AMPK) was significantly reduced in DAPIT OE cells, reflecting a state of metabolic saturation caused by their increased ATP production. Conversely, DAPIT KO cells exhibited a stress-induced arrest profile. These cells showed an upregulation of AMPK, which was accompanied by a significant transcriptional upregulation of the tumor suppressor *p53* (Figure 3E), which likely acts as the primary driver of the observed growth arrest, while *p16* and *p21* expression levels remained unchanged compared to wildtype cells. Interestingly, the expression of the key growth and metabolic regulator mechanistic target of rapamycin (mTOR) remained unchanged in both DAPIT OE and DAPIT KO cells (Figure 3E).

### Deficiency of DAPIT induces oxidative stress and triggers stress signaling

Our previous bioenergetic analyses revealed that DAPIT OE caused enhanced ATP production despite a reduced mitochondrial mass, suggesting that OXPHOS activity is more efficient and coupled in these cells. We hypothesized that DAPIT abundance directly influences mitochondrial reactive oxygen species (mtROS) levels and downstream cell survival. To test this, we performed live-cell imaging using the superoxide indicator MitoSOX normalized to mitochondrial mass via MTG staining. DAPIT OE cells exhibited decreased mtROS levels compared to wildtype cells (Figure 3F). This state was accompanied by a significant transcriptional upregulation of the antioxidant enzyme glutathione peroxidase 7 (*GPX7*) in DAPIT OE cells (Figure 3G), indicating an enhanced capacity to scavenge potential mtROS. In contrast, DAPIT KO cells exhibited a tendency toward elevated mtROS (Figure 3F). This oxidative stress in DAPIT KO cells was accompanied by a significant downregulation of *GPX7* (Figure 3G). This impairment of the enzymatic antioxidant shield highlights the vulnerability of DAPIT-deficient cells to oxidative damage. Given that these cellular profiles suggest divergent levels of mitochondrial health, we next examined whether these changes alter the rate of mitochondrial quality control via mitophagy. We quantified mitophagy by determining the colocalization of lysosomes (LysoTracker Red, LTR) with mitochondria (MTG). In DAPIT OE cells, we observed a significant reduction in colocalization (Figure 3H), suggesting decreased mitophagy. In DAPIT KO cells, mitophagy rates remained unchanged compared to wildtype cells, despite a reduction in mitochondrial mass. To investigate whether this may trigger apoptosis, we assessed the colocalization of cytochrome *c* (cyt *c*) with mitochondria (marked by the receptor subunit TOM20 of the translocase of the outer membrane) to evaluate mitochondrial outer membrane permeabilization. Both DAPIT OE and DAPIT KO cells showed a significant increase in cyt *c*/TOM20 colocalization compared to wildtype cells (Figure 3I and Figure S3D), indicating a reduction in cytochrome *c* release and a subsequent suppression of apoptosis. To understand this phenotype, we analyzed the transcriptional profiles of key apoptotic regulators. In DAPIT OE cells, the reduction in apoptotic signaling was reflected by a coordinated transcriptional downregulation of Bcl-2-associated X protein (*Bax*; pro-apoptotic pore former) and B-cell lymphoma 2 (*Bcl2*; anti-apoptotic regulator) genes, confirming a highly resilient, anti-apoptotic state (Figure 3J). The final blockade of the cascade is confirmed downstream by the significant reduction in caspase 9 (*Casp9*; initiator caspase) expression. In contrast, DAPIT KO cells showed significant transcriptional upregulation of pro-apoptotic *Bax* and anti-apoptotic *Bcl2*. In summary, DAPIT levels act as a metabolic switch that regulates cell survival. DAPIT OE cells achieve an extremely resilient state that requires only minimal quality control efforts. Conversely, DAPIT KO cells exhibit a bioenergetic bottleneck that makes them susceptible to oxidative stress.

### Quantitative proteomic profiling reveals structural and metabolic reprogramming in DAPIT overexpressing cells

Since accelerated cell proliferation is associated with an increased need for energy and metabolic precursors, rapidly dividing cells typically upregulate glycolysis. We therefore investigated the glycolytic activity by analyzing the intracellular glycolytic enzymes and intermediates. We first focused on hexokinase 2 (HK2), which catalyzes the first step of glycolysis. We used a coupled enzymatic assay to determine HK2 activity in fresh cell lysates (Figure 4A). DAPIT OE cells showed a significant increase in HK2 activity compared to wildtype cells, while DAPIT KO cells remained unchanged. This enzymatic upregulation was directly mirrored by an increased expression of the *HK2* gene in DAPIT OE cells (Figure 4B). The specificity of the coupled enzymatic assay was validated using the competitive inhibitor 2-deoxy-D-glucose (2-DG), which completely inhibited HK2 activity in all cell lines (Figure S4B). To determine whether the increased HK2 activity in DAPIT OE cells actually leads to an accumulation of downstream glycolytic metabolites, we indirectly measured endogenous pyruvate levels using a coupled lactate dehydrogenase assay (Figure 4C). Since the rate of NADH depletion in this assay is directly proportional to the amount of pyruvate present in the sample, we used this method to estimate pyruvate levels. DAPIT OE cells exhibited significantly faster NADH depletion, indicating increased cellular pyruvate levels, whereas DAPIT KO cells remained at the same level as wildtype cells (Figure 4C). The specificity of the assay was validated using measurements without cell lysates, which showed no depletion of NADH (Figure S4C). This pyruvate accumulation, combined with the increased HK2 activity, supports our model that DAPIT OE cells display accelerated glycolysis. To functionally assess glycolysis in living cells, we also performed a glycolysis stress test using a Seahorse XF96 Flux Analyzer to measure the extracellular acidification rate (ECAR). DAPIT OE cells exhibited a significantly reduced ECAR compared to wildtype cells (Figure S4D), while DAPIT KO cells showed no significant differences. This unexpected ECAR profile did not reflect the bioenergetic demands of rapid cell proliferation observed in DAPIT OE cells, but rather indicates that pyruvate is not converted to lactate but efficiently funneled into the mitochondria. In conclusion, our results show that DAPIT plays a central coordinating role in energy metabolism, and that its overexpression induces a hyperproliferative state in the cell, driven by a highly efficient, synchronized acceleration of both glycolysis and OXPHOS. Next, we determined global changes in metabolic and structural rewiring by proteomic analysis. Quantitative mass spectrometry (MS) revealed a striking difference in the extent of proteomic changes between the two DAPIT models. Analysis using a volcano plot demonstrated that DAPIT OE induced a widespread proteomic remodeling, which resulted in 324 proteins exhibiting significantly different expression levels compared to wildtype cells (Figure 4D). In contrast, the proteome of DAPIT KO cells remained largely unchanged and showed only two significant changes (Figure 4D). One of the downregulated proteins was DAPIT (ATP5MK) itself, which confirmed both the effectiveness of the genetic KO and the reliability of the MS dataset. The only other significantly downregulated protein in the DAPIT KO model was the tissue-nonspecific alkaline phosphatase (ALPL), which suggests a limited metabolic response rather than a comprehensive systemic adaptation. To systematically categorize the proteomic changes in DAPIT OE cells, we performed a Gene Ontology enrichment analysis. The most enriched terms were related to cellular connectivity (anchoring and cell junctions, cadherin binding) and the spatial architecture of the cytoskeleton (interaction with cytoskeletal proteins, activity of structural molecules), as well as with markers for vesicular transport (extracellular exosomes) (Figure 4E). These proteomic phenotypes are consistent with the morphological changes observed in DAPIT OE cells. To identify changes at the signaling pathway level, we performed a Gene-Set Enrichment Analysis (Figure 4F, S5A). “Collagen biosynthesis” and “mitochondrial translation” emerged as downregulated signaling pathways. A leading-edge analysis confirmed suppression of several collagen isoforms (COL) and mitochondrial ribosomal subunits (MRPL) (Figure S5C, D), consistent with the reduced mitochondrial mass in DAPIT OE cells. The positively enriched signaling pathways were associated with the cell cycle and cell division, particularly “kinesins” and “processes mediated by Polo-like kinases” (Figure 4F, S5B). This enrichment was driven by the upregulation of mitotic spindle motor proteins (KIF family) and key cell cycle regulators, including Polo-like kinase 1 (PLK1) and cyclins B1/B2 (CCNB1/B2) (Figure S5E, F). Overall, the proteome of DAPIT OE cells reflects a shift away from extracellular matrix production and mitochondrial biogenesis, thereby promoting the upregulation of the mitotic apparatus necessary for hyperproliferation.

**Figure 4:**
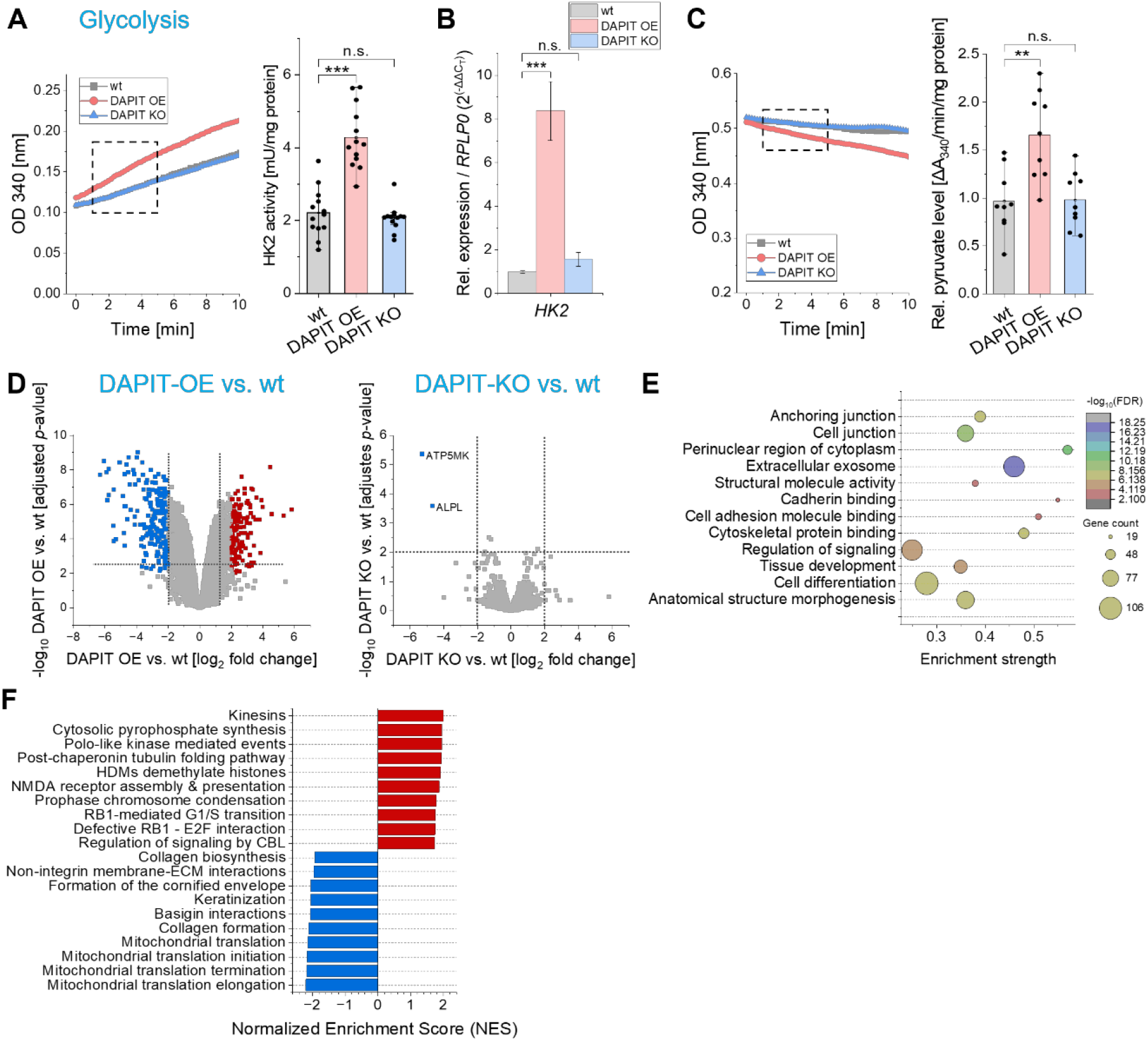
Metabolic rewiring initiated by DAPIT overexpression. **(A)** Representative kinetic profile of hexokinase 2 (HK2) activity from fresh cell lysates of HeLa wildtype (wt), DAPIT overexpressing (OE), and DAPIT knockout (KO) cells, monitored by the accumulation of NADH over time. The assay was coupled to an excess of exogenous glucose-6-phosphate dehydrogenase to continuously convert HK2-generated glucose-6-phosphate. Reduction of NAD^+^ to NADH was measured in real-time via absorbance (optical density, OD) at 340 nm using equal amounts of total protein (50 µg cell lysate). Quantification of HK2 activity was calculated from linear slope of kinetic curve (minutes 1-6, dashed frame) using the extinction coefficient of NADH (N=4, n=3 wells; Kruskal-Wallis with Dunn’s *post-hoc* test). **(B)** Quantification of relative (rel.) mRNA expression levels of *HK2*. Expression levels were determined by quantitative real-time PCR and normalized to housekeeping gene *RPLP0* using the 2^-ΔΔCt^ method with control group (wt) set to 1 (fold change) (N=5, n=3; one-way ANOVA with Scheffé’s *post-hoc* test). **(C)** Representative kinetic profile of endogenous pyruvate consumption from fresh cell lysates of HeLa wt, DAPIT OE, and DAPIT KO cells, monitored by the depletion of NADH. The assay was coupled to an excess of exogenous lactate dehydrogenase and NADH to continuously convert endogenous pyruvate into lactate. Oxidation of NADH to NAD^+^ was measured in real-time via the decrease in absorbance at 340 nm using equal amounts of total protein (50 µg cell lysate). Quantification of relative pyruvate levels was calculated from linear slope of the kinetic curve (minutes 1-6, dashed frame) using the extinction coefficient of NADH (N=3, n=3 wells; one-way ANOVA with Scheffé’s *post-hoc* test). **(D)** Volcano plots visualizing differentially expressed proteins in DAPIT OE cells versus HeLa wt cells, and DAPIT KO cells versus wt cells. X-axis represents the log_2_ FC and y-axis shows the **-**log_10_ (adjusted *p*-value). Dashed lines indicate applied significance thresholds (log2 FC≥2 and ≤**-**2, adjusted *p*-value ≤0.01). Highlighted points show significantly upregulated (red) and downregulated (blue) proteins. **(E)** Gene Ontology enrichment analysis (bubble plot) of significantly regulated proteins in DAPIT OE cells. Bubble size corresponds to number of genes within each term, and color scale reflects the significance level (FDR). **(F)** Gene Set Enrichment Analysis displaying top enriched pathways based on Reactome database. Bar plot shows the normalized enrichment score (NES) for significantly upregulated (red) and downregulated (blue) pathways in DAPIT OE cells. Data are presented as mean ± SD of N=3-5 independent biological experiments. The exact number of independent experiments (N) and each number of analyzed technical replicates, wells, or individual cells (n) are indicated in the respective brackets for each panel. Individual data points are overlaid in (A) and (B). Mass spectrometry data were obtained from N=4 independent biological experiments. Statistics: ***p≤0.001; **p≤0.01; *p≤0.05. (See also Figure S4 and S5). ANOVA, analysis of variance; DAPIT, diabetes-associated protein in insulin-sensitive tissue; FC, fold change; FDR, false discovery rate; NAD^+^, oxidized nicotinamide adenine dinucleotide; NADH, reduced nicotinamide adenine dinucleotide; RPLP0, ribosomal protein lateral stalk subunit P0; SD, standard deviation.

### DAPIT expression is upregulated during muscle differentiation and in insulin-sensitive tissues

Our findings demonstrated that DAPIT levels act as a metabolic switch in HeLa cells. Therefore, we next investigated what physiological relevance DAPIT expression could have across different cell types and metabolic states. Because DAPIT was originally identified as a protein expressed in insulin-sensitive tissues and is encoded by the *USMG5* gene [11], we compared its baseline protein abundance in HeLa wildtype cells with that in mouse skeletal muscle (C2C12) and mouse adipose (3T3-L1) cell lines (Figure 5A, B). Western blot analysis revealed that undifferentiated C2C12 myoblasts exhibited significantly higher relative protein levels of DAPIT compared to HeLa wildtype cells (Figure 5A). To determine whether DAPIT abundance is further modulated during cellular maturation, we also assessed relative DAPIT protein levels after differentiation. The transition from proliferating C2C12 myoblasts to multinucleated myotubes was accompanied by a significant upregulation of DAPIT protein levels. Similarly, upon induction of adipogenesis, differentiated 3T3-L1 adipocytes also displayed an increase in DAPIT protein levels compared to their pre-adipocyte precursors (Figure 5B). Taken together, these data show that DAPIT protein levels are not only upregulated during muscle differentiation but is also elevated in insulin-sensitive tissues.

**Figure 5:**
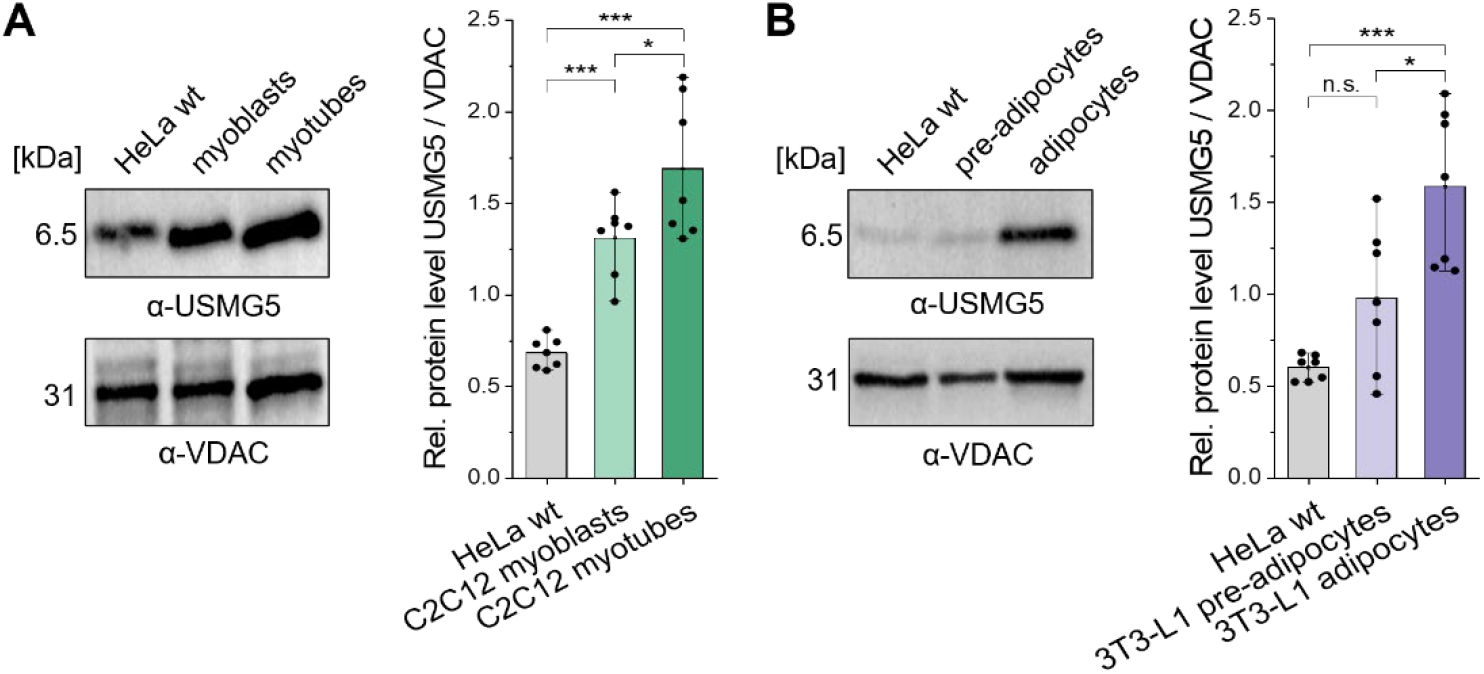
DAPIT levels are upregulated during cellular differentiation. **(A, B)** Representative Western blots and quantification of relative (rel.) protein levels of USMG5 (DAPIT) in **(A)** HeLa wildtype (wt), C2C12 mouse myoblasts, differentiated C2C12 mouse myotubes, and in **(B)** 3T3-L1 mouse pre-adipocytes and differentiated 3T3-L1 mouse adipocytes normalized to mitochondrial loading control VDAC (N=3, n=2; one-way ANOVA with Scheffé’s *post-hoc* test). Data are presented as mean ± SD from three independent biological experiments. The exact number of independent experiments (N) and each number of analyzed technical replicates, wells, or individual cells (n) are indicated in the respective brackets for each panel. Statistics: ***p≤0.001; **p≤0.01; *p≤0.05. ANOVA, analysis of variance; DAPIT, diabetes-associated protein in insulin-sensitive tissue; SD, standard deviation; USMG5, up-regulated during skeletal muscle growth protein 5; VDAC, voltage-dependent anion channel.

## DISCUSSION

Mitochondrial F_1_F_O_-ATP synthase is the most important enzyme for oxygenic cellular ATP production, but the regulatory role of its peripheral accessory subunits remains only partially understood. While electron microscopy studies have resolved the static architecture of the mammalian ATP synthase complex [13, 16], we here showed that the accessory subunit DAPIT (USMG5) acts as a crucial metabolic switch that directly links the oligomerization and stabilization of ATP synthase to the regulation of cellular bioenergetics and proliferative capacity of the cell.

A key physiological paradox identified in our study is the inverse relationship between mitochondrial mass and bioenergetic performance induced by DAPIT overexpression (OE). Although DAPIT OE cells exhibit a significant reduction in mitochondrial mass and a downregulation of mitochondrial ribosomal subunits (Figure 3, 4), they achieve a state of metabolic hyper-efficiency. This state is characterized by increased oxygen consumption, enhanced ATP production, and a hyperpolarized ΔΨ_m_ (Figure 2). The saturation of DAPIT binding sites forces ATP synthase into highly stable dimeric configurations (Figure 1). It is well established that the dimerization of ATP synthase is the fundamental prerequisite for generating the high membrane curvature at the cristae tips [6, 7]. Our data complement these findings by identifying DAPIT as an active driver of this ATP synthase stabilization. Stabilization of ATP synthase oligomers likely creates a tightly sealed proton sink at the tips of the cristae, minimizing nonspecific proton leakage. This tight coupling is further evidenced by the significant downregulation of the uncoupling proteins *UCP2* and *UCP3* in DAPIT OE cells (Figure 2). In contrast, DAPIT KO cells exhibit an upregulation of *UCP2*, which likely acts as an active bypass for protons to mitigate the stress caused by the structural destabilization of the ATP synthase oligomers. Despite the upregulation of *UCP2*, we made the paradoxical observation that DAPIT KO cells maintain a stable ΔΨ_m_. A classic compensatory mechanism to sustain ΔΨ_m_ during structural dysfunction or uncoupling is the reverse function of the ATP synthase, which then consumes ATP to pump protons back into the intermembrane space. However, our data exclude this reverse ATPase activity in DAPIT KO cells. First, their total ATP hydrolysis capacity was not increased (Figure 2A). Second, if reverse ATPase activity were responsible for sustaining the ΔΨ_m_, treatment with the ATP synthase inhibitor oligomycin would trigger a depolarization. Instead, the ΔΨ_m_ of DAPIT KO cells remained insensitive to oligomycin, indicating that their stable membrane potential is maintained independently of ATP hydrolysis (Figure 2).

To fuel the dynamic upregulation of OXPHOS, DAPIT OE cells require an accelerated influx of substrates. Rapidly proliferating cells typically rely on aerobic glycolysis (the Warburg effect), which is usually characterized by high extracellular acidification rates (ECAR) due to lactate secretion [30]. However, our metabolic flux analysis revealed a decreased ECAR in DAPIT OE cells (Figure S4). Therefore, we hypothesized a spatial “funnel effect”. It is well established that HK2 physically localizes to the outer mitochondrial membrane via VDAC to directly couple glycolysis to mitochondrial ATP production [31]. While the overall extracellular glycolytic flux appeared to remain low, the specific activity of hexokinase 2 (HK2) was significantly upregulated, leading to an increase in intracellular pyruvate levels (Figure 4). This synthesized pyruvate is likely directed into the hyperactive mitochondria to fuel the tricarboxylic acid cycle and OXPHOS, thereby bypassing its conversion to lactate. Consequently, intracellular pyruvate levels are increased in the steady-state. Together, this demonstrates a highly synchronized acceleration of glycolysis and OXPHOS, which drives the cell’s hyperproliferative state.

The increase in metabolic activity leads to a redistribution of cellular resources, as evidenced by quantitative proteome profiling (Figure 4). By utilizing an efficient, stabilized pool of active ATP synthases, DAPIT OE cells are able to suppress energy-intensive, non-essential processes such as the production of the extracellular matrix (collagens) and mitochondrial translation. Instead, the energy conserved from downregulating these processes is redirected to support the mitotic apparatus, particularly the spindle motor proteins (kinesins) and Polo-like kinase 1 (PLK1). This molecular reprogramming directly translates into accelerated proliferation rates and the absence of AMPK-mediated energy stress observed *in vitro* (Figure 3). Furthermore, the better coupling of electron transport chain to ATP phosphorylation protects the cell from oxidative damage. The continuous, efficient flow of electrons prevents the accumulation of mtROS (Figure 3) and thereby eliminates the need for the constant, energy-intensive transcriptional upregulation of apoptotic regulators (*Bax*/*Bcl2*). In contrast, structural inefficiencies in DAPIT KO cells make them vulnerable and reliant on compensatory survival mechanisms.

Finally, our data show that DAPIT-mediated bioenergetic optimization is not an isolated phenomenon of an immortalized cancer cell line, but rather reflects a highly conserved physiological mechanism. The significant upregulation of DAPIT protein levels during the differentiation of C2C12 myoblasts into myotubes and of 3T3-L1 preadipocytes into mature adipocytes (Figure 5) is consistent with the metabolic transition required in these tissues. As these cells transition from a proliferating, glycolytic precursor state to a mature, insulin-sensitive state, they require highly efficient OXPHOS to maintain muscle contraction and lipid metabolism [8, 32]. This dynamic assembly of DAPIT to the ATP synthase is structurally feasible, since DAPIT is incorporated in the last step of ATP synthase biogenesis and is not required for the assembly of other subunits [16]. Its significance is further demonstrated by the fact that mutations in the *USMG5* gene result in Leigh syndrome, a severe and progressive neurodevelopmental disorder linked to mitochondrial energy deficiency [11].

In summary, we propose that the incorporation of DAPIT functions as an adjustable structural anchor for the ATP synthase. While basal DAPIT concentrations are sufficient to maintain basic monomeric functionality, its targeted upregulation stabilizes higher-order oligomers and places the IMM in a state of maximum bioenergetic efficiency. Our results show that DAPIT acts as a central metabolic switch that translates the architecture of the ATP synthase into cellular resilience and hyperproliferation, thereby underscoring the significance of supramolecular organization of the OXPHOS system – and in particular ATP synthase – as a fundamental determinant of cell metabolism.

### Limitations of the study

The study investigated DAPIT effects in a cancer cell line that is partially polyploid. The advantage of this cell model is its metabolic flexibility, characterized by the Warburg effect; but it is also its limitation, as this mixed metabolism is typical for cancer cells and likely differs from stem and differentiated cells. Therefore, future studies should build on our observation that DAPIT is upregulated during adipocyte and myoblast differentiation to further decipher its role in metabolic switches during cellular differentiation and tissue development.

## RESOURCE AVAILABILITY

### Lead contact

Requests for further information and resources should be directed to and will be fulfilled by the lead contact, Karin B. Busch.

### Materials availability

Cell lines and plasmids generated in this study are available upon request from the lead contact.

### Data and code availability

• LC-MS/MS raw data and processed data are available at ProteomeXchange: PXD087595. Microscopy data reported in this paper will be shared by the lead contact upon request.

• This paper does not report original code.

• Any additional information required to reanalyze the data reported in this paper is available from the lead contact upon request.

## Supporting information

Manuscript

## ACKNOWLEDGEMENTS

We thank Frank Schmelter for valuable technical support.

## AUTHORS CONTRIBUTIONS

M.T.B and K.B.B. designed the experiments; M.T.B and P.M.O. performed the experiments and analyzed the data. M.S. and M.H. did mass spectrometry analysis and provided results. M.T.B. and K.B.B. wrote the manuscript. K.B.B. supervised the project and secured funding. All authors read and approved the manuscript.

## DECLARATION OF GENERATIVE AI AND AI-ASSISTED TECHNOLOGIES IN THE WRITING PROCESS

During the preparation of this work, the authors used ChatGPT 5.2 (OpenAI)/DEEPL to improve the language clarity of the manuscript. After using this tool, the authors reviewed and edited the content as needed and take full responsibility for the content of the published article.

## STAR * METHODS

Detailed methods are provided in the online version of this paper and include the following:

- KEY RESOURCES TABLE
- EXPERIMENTAL MODEL

`o` Mammalian cell culture
- METHOD DETAILS

`o` Plasmid Construction
`o` Transfection of mammalian cells
`o` Cell proliferation kinetics
`o` Quantitative real-time polymerase chain reaction
`o` Mitochondrial isolation and protein quantification
`o` SDS-PAGE and immunoblotting
`o` Protein turnover and half-life determination
`o` Native PAGE
`o` In-gel ATP hydrolysis activity assay
`o` Fluorescence microscopy and live-cell imaging
`o` Mitochondrial membrane potential (ΔΨ_m_)
`o` Mitochondrial mass and cellular morphology
`o` Mitochondrial ROS measurements
`o` Mitochondrial and lysosomal colocalization
`o` Cytochrome c release assay
`o` Real-time ATP production rate analysis
`o` Luciferase-based mitochondrial ATP synthesis assay
`o` Mitochondrial respiration analysis
`o` Glycolytic function analysis (Glycolysis Stress Test)
`o` Hexokinase 2 enzymatic activity assay
`o` Intracellular pyruvate quantification
`o` Proteomic sample preparation and LC-MS/MS analysis
`o` Data processing and bioinformatics
- QUANTIFICATION AND STATISTICAL ANALYSIS

## SUPPLEMENTAL INFORMATION

Supplemental Information includes five figures (S1-S5) and three tables and can be found with this article online.

## STAR * METHODS

### KEY RESOURCES TABLE

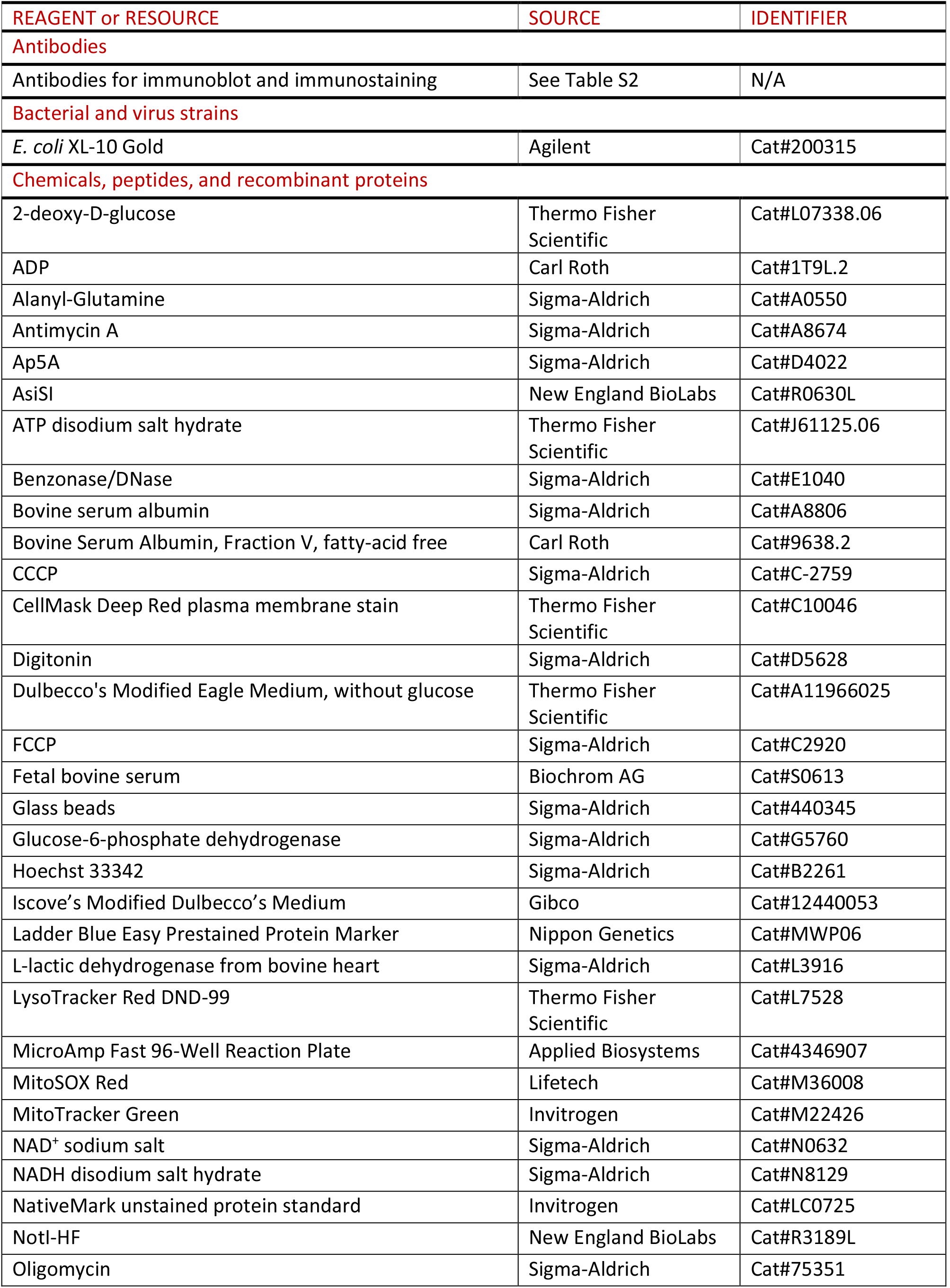
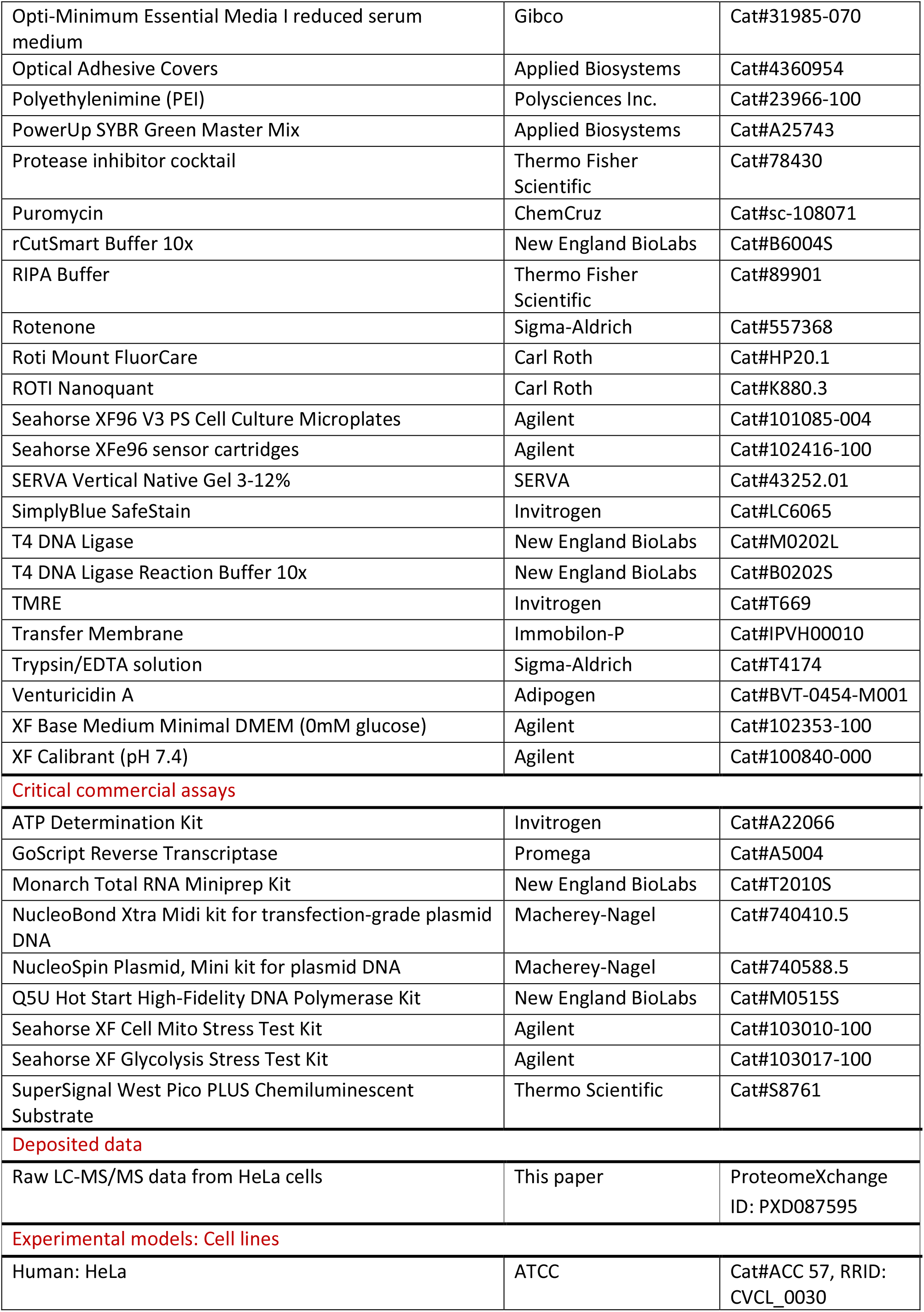
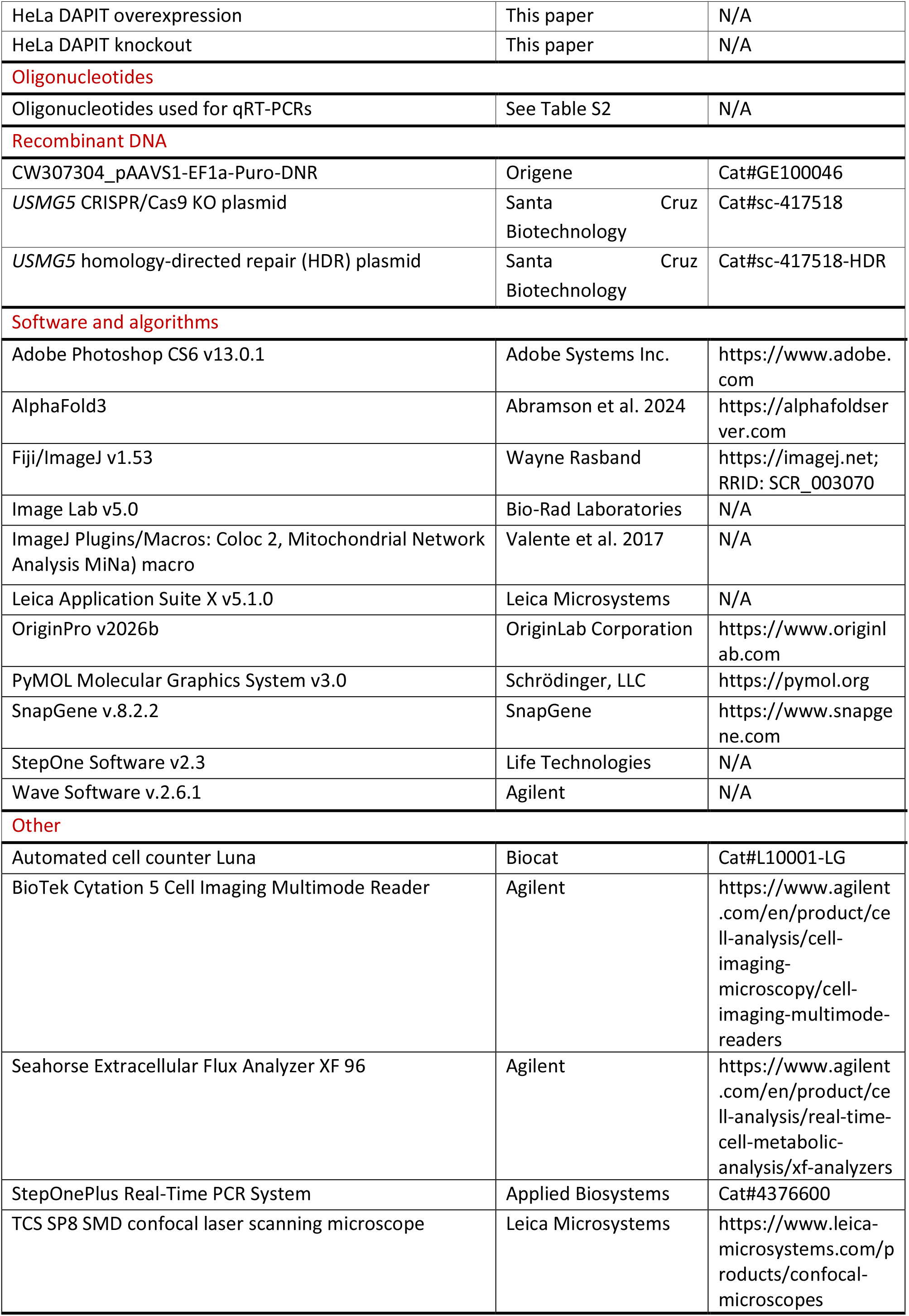

### EXPERIMENTAL MODEL AND STUDY PARTICIPANT DETAILS

#### Mammalian cell culture

HeLa cells were purchased from the Leibnitz Institute DMSZ [#ACC 57] and cultured in high-glucose Iscove’s Modified Dulbecco’s Medium (25 mM glucose) supplemented with 10% fetal bovine serum FBS and 1% penicillin-streptomycin at 37°C and 5% CO_2_. Cells were maintained at 70-80% confluence and passaged every 2-3 days using standard trypsin-EDTA detachment. To induce an oxidative phosphorylation (OXPHOS)-dependent metabolic shift, cells were adapted for a minimum of four weeks in glucose-free galactose medium (Dulbecco’s Modified Eagle Medium, DMEM, supplemented with 10% FBS, 5.5 mM galactose, 1% HEPES, 1% MEM non-essential amino acids, 1 mM pyruvate, and 1% penicillin-streptomycin). Murine C2C12 myoblasts were cultured in high-glucose Dulbecco’s Modified Eagle Medium (DMEM, 25 mM glucose) supplemented with 10% FBS and 1% penicillin-streptomycin at 37°C in a humidified incubator at 5% CO_2_. To induce myogenic differentiation, cells at 80-90% confluence were shifted to a low-serum medium (DMEM, 2% horse serum, 1% penicillin-streptomycin), which was exchanged every 48 h for seven days until multinucleated myotubes formed. Murine 3T3-L1 pre-adipocytes were maintained in high-glucose DMEM containing 10% newborn calf serum (NCS), 2 mM L-glutamine, and 1% penicillin-streptomycin at 37°C and 10% CO_2_. Adipogenesis was initiated at 80-90% confluence using a three-step protocol. Briefly, cells were stimulated for three days in high-glucose DMEM supplemented with 10% FBS, 1% penicillin-streptomycin, 2 mM alanyl-glutamine (Ala-Gln), 0.5 mM 3-isobutyl-1-methylxanthine, 1 µM insulin, 0.25 µM dexamethasone, and 1 µM troglitazone. From days 3 to 6, IBMX and dexamethasone were omitted from the medium. From day 6 onwards, differentiating adipocytes were maintained in basal medium (high-glucose DMEM, 10% FBS, 1% penicillin-streptomycin, 2 mM Ala-Gln), which was refreshed every 2-3 days until intracellular lipid accumulation was visually confirmed.

### METHOD DETAILS

#### Plasmid Construction

To generate DAPIT expression vector, the *USMG5* coding sequence was amplified from wildtype HeLa cDNA using Q5U Hot Start High-Fidelity DNA Polymerase Kit (New England BioLabs M0515S) and cloned into the AsiSI (New England BioLabs R0630L) and NotI (New England BioLabs R3189L) restriction sites of the CW307304_pAAVS1-EF1a-Puro-DNR (Origene GE100046) backbone using T4 DNA ligase (New England BioLabs M0202L). Plasmids were transformed into competent XL10-Gold *E. coli* (Agilent 200315) using standard heat-shock procedures. All constructs were purified using the NucleoBond PC 100 Midi kit (Macherey-Nagel, 740410.50) and verified by Sanger sequencing prior to mammalian transfection.

#### Transfection of mammalian cells

To generate stable DAPIT overexpression (OE) and DAPIT knockout (KO) models in HeLa cells, a CRISPR/Cas9-mediated approach was used. Cells were co-transfected with a *USMG5* CRISPR/Cas9 KO plasmid and a corresponding homology-directed repair (HDR) plasmid (Santa Cruz Biotechnology sc-417518, sc-417518-HDR) at an equimolar ratio utilizing polyethylenimine (PEI; 25 kDa; Polysciences 23966-100) at a 2:1 PEI to DNA ratio. Following a 24 h recovery period, successfully transfected cells were selected in culture medium supplemented with 1 µg/ml puromycin. To ensure genetic homogeneity, the selected stable pools of DAPIT KO were subjected to limiting dilution in 96-well microplates. After visually confirming that the cells originated from a single colony, a monoclonal cell line was isolated and clonally expanded for all subsequent experiments.

#### Cell proliferation kinetics

To determine cellular proliferation kinetics, HeLa cells were seeded in triplicate into 12-well plates at a density of 10,000 cells per well. At 24, 48, 72, and 96 h post-seeding, nuclei were stained with Hoechst 33342 (2 µg/ml; Sigma-Aldrich B2261) for 10 min. Automated image acquisition and quantitative nuclear segmentation were performed utilizing a Cytation 5 Cell Imaging Multimode Reader (Agilent). To account for initial seeding variations, absolute cell counts were normalized to the 24 h baseline measurement. Finally, specific proliferation rates were calculated from the slope of the exponential growth phase.

#### Quantitative real-time polymerase chain reaction

Total RNA was isolated utilizing the Monarch Total RNA Miniprep Kit (New England BioLabs T2010S) and quantified spectrophotometrically. Complementary DNA was synthesized utilizing the GoScript Reverse Transcriptase Kit (Promega A5004) with random hexamer primers. Quantitative real-time PCR was performed on a StepOnePlus Real-Time PCR System (Applied Biosystems 4376600) using PowerUp SYBR Green Master Mix (Applied Biosystems A25743) with a final primer concentration of 200 nM. The thermal cycling conditions included an initial denaturation at 95°C for 10 min, followed by 40 cycles of 95°C for 15 s and 60°C for 60 s, concluding with a melting curve analysis to confirm product specificity. Primer sequences (Eurofins Genomics) are provided in Supplementary Table 1. Relative mRNA expression was calculated according to the 2^-ΔΔCt^ method [33], normalized to the internal housekeeping gene *RPLP0*, and expressed as fold change relative to wildtype cells.

#### Mitochondrial isolation and protein quantification

Mitochondrial fractions were isolated applying a standard differential centrifugation protocol at 4°C. Briefly, harvested cells were resuspended in ice-cold isolation buffer (1 mM EGTA, 5 mM HEPES, 210 mM mannitol, 70 mM sucrose) and mechanically disrupted utilizing a tissue homogenizer. Unbroken cells and nuclei were pelleted by repeated centrifugation at 800 × g for 10 min. The pooled supernatants were subsequently clarified at 1000 × g for 10 min, followed by mitochondrial pelleting at 10,000 × g for 10 min. The resulting mitochondrial pellets were resuspended in isolation buffer. Protein concentrations of total cell lysates and isolated mitochondria were determined utilizing the ROTI Nanoquant assay (Carl Roth, K880.3). Absorbance ratios at 590 nm and 450 nm were acquired on a Cytation 5 Multimode Reader (Agilent) and absolute concentrations were calculated via linear regression against a bovine serum albumin (BSA) standard curve.

#### SDS-PAGE and immunoblotting

Cells were lysed in RIPA buffer (Thermo Fisher Scientific 89901) supplemented with protease inhibitor cocktail (1:100, Thermo Fisher Scientific 78430), and benzonase/DNase (1:1000, Sigma-Aldrich E1040). Following a 20-min incubation on ice, lysates were cleared, and protein concentrations were determined via Bradford assay. Equal amounts of total protein (25 µg) were mixed with 4× LDS sample buffer (30 mM Tris pH 6.8, 0.05% Bromo Phenol Blue, 34.8% glycerol, 40% LDS, 4% β-mercaptoethanol), denatured at 90°C for 5 min, and resolved on 12% SDS-polyacrylamide gels. Proteins were wet-transferred onto methanol-activated PVDF membranes (Immobilon-P IPVH00010) at 25 V overnight at 4°C. Membranes were blocked with either 10% non-fat milk or 5% BSA in TBST for 1 h at room temperature (RT) and probed with primary antibodies (Supplementary Table 2) overnight at 4°C. After three TBST washes, membranes were incubated with HRP-conjugated secondary antibodies (1:2000) for 1 h at RT. Signals were detected using SuperSignal West Pico PLUS chemiluminescent substrate (Thermo Fisher Scientific S8761) on a BioRad Universal Hood III. Total protein levels were visualized by Coomassie staining. Quantification (integrated density) was performed using Fiji/ImageJ (v1.53), and target protein signals were normalized to the respective loading controls (GAPDH, VDAC, or total protein).

#### Protein turnover and half-life determination

Protein stability and turnover rates of ATP synthase and OPA1 were evaluated using a cycloheximide (CHX) chase assay. Briefly, cells were seeded in T25 flasks, allowed to adhere overnight, and subsequently treated with CHX (100 µg/ml) under standard culture conditions. The culture medium was replaced every 24 h. Cells were harvested at specified time points (0, 8, 24, 48, 72, and 96 h), lysed, and subjected to immunoblotting as described above. The relative protein abundance at 0 h was set to 100%. To determine degradation kinetics, relative signal intensities were plotted over time, and protein half-lives (t_1/2_) were calculated by linear regression analysis of the log-transformed data.

#### Native PAGE

Native protein assemblies were analyzed using clear native (CN) PAGE. Isolated mitochondria (50 µg) were solubilized on ice for 15 min in a buffer containing 50 mM imidazole, 50 mM NaCl, 2 mM 6-aminohexanoic acid, 1 mM EDTA, and 2% (w/v) digitonin, pH 7.0. Following centrifugation at >20,000 × g for 20 min at 4°C, the resulting supernatant was supplemented with buffer containing 0.1% Ponceau and 22.73% glycerol, and separated on a 3–12% SERVA Vertical Native Gel (43252.01). Electrophoresis was initiated at 80 V for 1 h at 4°C, with the voltage subsequently increased by 10 V/h until the dye front reached the bottom of the gel. Gels were stained with SimplyBlue SafeStain (Invitrogen LC6065) for 1 h to visualize total protein patterns and molecular weight markers, followed by destaining in ddH₂O. Proteins were wet-transferred onto PVDF membranes overnight as described above. To fix the proteins, membranes were incubated in 8% acetic acid for 15 min and washed with ddH₂O. Following air-drying, native marker bands were annotated, and residual Coomassie dye was removed by a 5-min methanol wash. After final washes in ddH₂O and TBST, immunoblotting and chemiluminescence detection were performed.

#### In-gel ATP hydrolysis activity assay

ATP hydrolysis capacity of the ATP synthase was assessed using an in-gel activity assay adapted from Wittig *et al.* [22]. Isolated mitochondria (100 µg) were solubilized, cleared by centrifugation (>20,000×g, 20 min, 4°C), and the resulting supernatant was mixed with Ponceau buffer, as described above for native PAGEs. Protein complexes were resolved on a 3–12% SERVA Vertical Native Gel (43252.01) at 4°C, with electrophoresis initiated at 80 V for 1 h and subsequently increased by 10 V/h until completion. Gels were pre-incubated in assay buffer (270 mM glycine, 35 mM Tris, pH 8.4) for 2 h at RT. For the enzymatic reaction, gels were transferred to a reaction buffer containing 270 mM glycine, 35 mM Tris, 4 mM ATP, 14 mM MgSO₄, and 0.2% Pb(NO₃)₂ (pH 8.4) and incubated overnight until white lead(II) phosphate precipitates became visible. The reaction was terminated by incubating the gel in 50% methanol for 30 min. To account for total protein loading, gels were subsequently stained with SimplyBlue SafeStain (Invitrogen LC6065) for 1 h and destained in ddH₂O for 2 h. Quantification of the precipitates was performed using Fiji/ImageJ (v1.53). ATPase activity was initially normalized to total protein input based on the Coomassie staining. To evaluate specific enzymatic activity independent of the absolute ATP synthase expression, this normalized activity was further adjusted to the relative ATP synthase protein abundance determined via corresponding SDS-PAGE analyses (with wildtype levels set to 1.0).

#### Fluorescence microscopy and live-cell imaging

All imaging was performed using a Leica TCS SP8 SMD confocal laser scanning microscope (cLSM) equipped with a 63× water objective (HC PL APO 63×/1.20 W motCorr CS2), a tunable white light laser (WLL), a 405 nm diode laser, and hybrid detectors (HyDs) with gallium arsenide phosphide (GaAsP) photocathodes. Live-cell imaging was conducted in a temperature- and CO_2_-controlled chamber (37°C, 5% CO2). Cells (20,000–30,000) were seeded into 8-well dishes 24 h prior to experiments. Image processing and quantitative analyses were performed using Fiji/ImageJ (v1.53).

#### Mitochondrial membrane potential (ΔΨ_m_)

To assess ΔΨ_m_, cells were co-stained with 7 nM TMRE (Invitrogen T669) and 100 nM MitoTracker Green (MTG; Invitrogen M22426) for 30 min at 37°C. Following three washes, cells were imaged in medium containing 7 nM TMRE. Where indicated, cells were pre-treated with oligomycin (5 µM, 30 min), venturicidin A (5 µM, 30 min), or CCCP (10 µM, 5 min). MTG and TMRE were excited at 488 nm and 554 nm, with emissions recorded at 498-549 nm and 564-746 nm, respectively. Ratiometric analysis (TMRE/MTG integrated density per cell) was performed following automated background subtraction setting the background to NaN.

#### Mitochondrial mass and cellular morphology

For mitochondrial mass and cell area quantification, live cells were co-stained with 100 nM MTG (30 min) and 5 µg/ml CellMask Deep-Red (Thermo Fisher Scientific C10046; 10 min) at 37°C. MTG (λex=488 nm; λem=498-549 nm) and CellMask (λex=641 nm; λem=660-779 nm) signals were acquired. Mitochondrial mass was quantified utilizing the Mitochondrial Network Analysis (MiNa) toolset [23], extracting the mitochondrial footprint from skeletonized binary masks. Single-cell mitochondrial mass was normalized to the respective total cellular area determined via CellMask boundary segmentation. For nuclear morphometry, cells were labeled with 1 µg/ml Hoechst 33342 (Sigma-Aldrich B2261; λex=405 nm, λem=415-605 nm) and CellMask Deep-Red. Nuclear area was extracted via automated thresholding of the Hoechst signal.

#### Mitochondrial ROS measurements

Mitochondrial superoxide levels were evaluated by co-staining cells with 2.5 µM MitoSOX Red (Lifetech M36008) and 100 nM MTG for 30 min at 37°C. Cells were washed and imaged in medium containing 2.5 µM MitoSOX Red (λex=561 nm; λem=660–779 nm). Ratiometric quantification (MitoSOX/MTG) was performed as described for ΔΨ_m_ measurements.

#### Mitochondrial and lysosomal colocalization

Mitophagy was assessed via live-cell dual-fluorescence imaging of mitochondria and lysosomes. Cells were stained with 100 nM LysoTracker Red DND-99 (LTR; Thermo Fisher Scientific L7528) for 1 h and 100 nM MTG for 30 min at 37°C. LTR was excited at 573 nm (λem=583-750 nm). Spatial colocalization of the lysosomal and mitochondrial networks was quantified on a single-cell basis by calculating the Manders’ overlap coefficient using the Coloc 2 plugin in Fiji.

#### Cytochrome c release assay

To evaluate mitochondrial outer membrane permeabilization, cells were fixed in 4% PFA (15 min, RT), permeabilized with 0.1% Triton X-100 (10 min), and blocked in 0.5% BSA/PBS (30 min). Cells were incubated with primary antibodies against cytochrome c and the mitochondrial marker TOM20 (Supplementary Table 2) for 1 h at RT, followed by corresponding Alexa Fluor-conjugated secondary antibodies (488 and 647; 1 h, RT, dark). Nuclei were counterstained with Roti Mount FluorCare DAPI (Carl Roth HP20.1). Cytochrome c release was quantified by calculating the Manders’ overlap coefficient between the cytochrome c and TOM20 signals using the Coloc 2 plugin in Fiji.

#### Real-time ATP production rate analysis

Real-time mitochondrial (mitoATP) and glycolytic (glycoATP) ATP production rates were quantified using a Seahorse XFe96 Flux Analyzer (Agilent). Cells were seeded at 25,000 cells/well in XF96 microplates and allowed to adhere overnight. Prior to the assay, the culture medium was replaced with unbuffered XF Minimal DMEM (pH 7.4, Agilent) supplemented with 25 mM D-glucose, 1 mM sodium pyruvate, and 2 mM L-glutamine. Cells were then incubated for 1 h at 37°C in a non-CO₂ incubator to ensure pH and temperature equilibration. Oxygen consumption rates (OCR) and extracellular acidification rates (ECAR) were recorded under baseline conditions and following the sequential injection of oligomycin (1.5 µM) and a rotenone/antimycin A mixture (0.5 µM each). The mitoATP and glycoATP production rates were calculated based on OCR, ECAR, and proton efflux rates (PER) using the Seahorse XF Real-Time ATP Rate Assay Report Generator within the Wave software (v2.6.1, Agilent). To account for variations in mitochondrial content between cell lines, the calculated metabolic rates were normalized to the mitochondrial mass, determined via MitoTracker Green fluorescence measurements (with wildtype levels set to 1.0).

#### Luciferase-based mitochondrial ATP synthesis assay

The functional ATP synthesis capacity of isolated mitochondria was quantified using a bioluminescence-based ATP Determination Kit (Invitrogen A22066). Freshly isolated mitochondria (50 µg protein, determined via Bradford assay) were suspended in 160 µl of mitochondrial respiration buffer (0.1% BSA fatty-acid free, 2 mM EDTA, 150 mM KCl, 10 mM KH_2_PO_4_, 0.1 mM MgCl_2_, 25 mM This-HCl). ATP production was fueled via Complex II by the addition of 5 mM succinate, with Complex I inhibited by 5 mM rotenone. The reaction mixture was further supplemented with 4 mM ADP as substrate and 20 µM Ap5A to inhibit adenylate kinase activity. The bioluminescent reaction was initiated by adding 10 µl of an enzyme-substrate mix (0.8 mM D-luciferin and 5 µg/ml firefly luciferase in 0.5 M Tris-acetate, pH 7.75). Luminescence was continuously recorded at 10-s intervals at 25°C using a Cytation 5 Cell Imaging Multimode Reader (Agilent). The basal ATP production rate was calculated from the linear slope of the luminescence increase between minutes 5 and 15. To isolate specific ATP synthase activity, the non-specific slope recorded in the presence of oligomycin (10 µM) was subtracted from the basal slope. Final oligomycin-sensitive ATP synthesis rates were normalized to mitochondrial protein input.

#### Mitochondrial respiration analysis

Mitochondrial respiratory function was evaluated using the Seahorse XF Cell Mito Stress Test on an XFe96 Flux Analyzer (Agilent). Cells were seeded at a density of 25,000 cells/well in XF96 microplates and allowed to adhere overnight. On the day of the assay, the culture medium was replaced with unbuffered XF Minimal DMEM (pH 7.4, Agilent) supplemented with 25 mM D-glucose, 1 mM sodium pyruvate, and 2 mM L-glutamine. The microplates were subsequently incubated for 1 h at 37°C in a non-CO₂ incubator for pH and temperature equilibration. Oxygen consumption rates (OCR) were recorded under baseline conditions and following the sequential injection of oligomycin (1 µM), FCCP (1 µM), and a rotenone/antimycin A mixture (1 µM each). Data acquisition and analysis were performed using Wave software (v2.6.1, Agilent). To account for variations in mitochondrial content between cell lines, OCR values were normalized to the mitochondrial mass, which was determined via MitoTracker Green fluorescence measurements (with wildtype levels set to 1.0).

#### Glycolytic function analysis (Glycolysis Stress Test)

Glycolytic function was assessed by measuring the extracellular acidification rate (ECAR) using the Seahorse XF Glycolysis Stress Test on an XFe96 Flux Analyzer (Agilent). Cells were seeded at a density of 25,000 cells/well in XF96 microplates and allowed to adhere overnight. Prior to the measurement, the culture medium was replaced with unbuffered XF Minimal DMEM (pH 7.4, Agilent) supplemented solely with 2 mM L-glutamine. The microplates were subsequently incubated for 1 h at 37°C in a non-CO₂ incubator. ECAR was recorded under baseline conditions and following the sequential injection of D-glucose (10 mM), oligomycin (1 µM), and 2-deoxy-D-glucose (50 mM). Data were acquired and analyzed using Wave software (v2.6.1, Agilent). To normalize ECAR values to cell number, cells were stained with Hoechst 33342 (2 µg/ml, Sigma-Aldrich) immediately following the final injection, and nuclei were counted using a Cytation 5 Cell Imaging Multimode Reader (Agilent).

#### Hexokinase 2 enzymatic activity assay

Hexokinase 2 (HK2) activity was determined using a coupled enzymatic assay. Cells were lysed on ice for 30 min, and lysates were cleared by centrifugation (15,000 × g, 10 min, 4°C). Protein concentrations were quantified via Bradford assay. Equal amounts of protein lysate (50 µg) were combined with reaction buffer to yield final assay concentrations of 50 mM Tris-HCl (pH 7.4), 10 mM MgCl₂, 25 mM D-glucose, 2 mM ATP, 0.5 mM NAD⁺, and 1 U glucose-6-phosphate dehydrogenase (Sigma-Aldrich G5760). The reduction of NAD⁺ to NADH was continuously monitored by measuring absorbance at 340 nm every 10 s for 30 min at 30°C using a Cytation 5 Cell Imaging Multimode Reader (Agilent). Specific HK2 activity was calculated from the linear slope of the reaction kinetics (minutes 1-6) using the NADH extinction coefficient, and normalized to total protein input. Assay specificity was confirmed via parallel control reactions utilizing 150 mM 2-deoxy-D-glucose to competitively inhibit HK2.

#### Intracellular pyruvate quantification

Relative intracellular pyruvate levels were evaluated using a kinetic spectrophotometric assay coupled to the lactate dehydrogenase (LDH)-catalyzed oxidation of NADH. Cell lysates were prepared and cleared as described above for the HK2 activity assay. Equal amounts of protein lysate (50 µg) were mixed with reaction buffer containing 50 mM Tris-HCl (pH 7.4), 10 mM MgCl₂, 25 mM D-glucose, 2 mM ATP, 0.4 mM NADH, and 2 U exogenous LDH (Sigma-Aldrich L3916). The rate of NADH consumption, which is directly proportional to the endogenous pyruvate level, was monitored by tracking the decrease in absorbance at 340 nm every 10 s for 30 min at 30°C on a Cytation 5 Cell Imaging Multimode Reader (Agilent). Relative pyruvate levels were calculated from the linear slope of NADH oxidation (minutes 1-6) using the NADH extinction coefficient, and normalized to total protein input. Assay specificity were verified using parallel control reactions omitting cell lysates.

#### Proteomic sample preparation and LC-MS/MS analysis

Cells were lysed in lysis buffer (10% glycerol, 150 mM NaCl, 50 mM Tris-HCl pH 7.4, 0.5% Triton X-100, protease inhibitor 1:100) supplemented with 0.5% sodium deoxycholate (SDC). Protein concentrations were adjusted to 2.5 µg/µl using lysis buffer and 25 µg (10 µl) of protein per sample was processed using a modified SP4 protocol [34]. Briefly, proteins were simultaneously reduced by addition of 1 µl of 100 mM tris(2-carboxyethyl)phosphine and alkylated with 1 µl of 400 mM chloroacetamide for 45 min at RT in the dark. Proteins were then precipitated onto glass beads (Sigma-Aldrich 440345; 48 µL of a 5 mg/mL suspension in acetonitrile, final acetonitrile concentration 80 %). Protein-bead complexes were washed twice with 100 µl 80% ethanol and digested overnight at 37°C with sequencing-grade trypsin (Promega, enzyme-to-protein ratio 1:50) in 80 µl of 50 mM ammonium bicarbonate containing 0.5% SDC. Following digestion, SDC was precipitated by acidification with 10% trifluoroacetic acid (pH<4). Cleared peptides were quantified using a DS-11 μVolume Spectrophotometer (DeNovix). LC-MS/MS analysis was performed on a system composed of a proteoElute nanoLC system (Bruker Daltonics) coupled online via a CaptiveSpray 2 source (capillary voltage 1600 V; dry gas 3.0 L/min at 200 °C) to a timsUltra AIP mass spectrometer (Bruker Daltonics). The peptide mixture (25 ng peptides in 1 µl 2% (v/v) acetonitrile in LC-MS grade water) was loaded directly onto a PepSep Advanced analytical column (25 cm × 75 µm, 1.5 µm particle size; Bruker Daltonics) at a pressure of 800 bar. Mobile phase A was 0.1% (v/v) formic acid in LC-MS grade water, and mobile phase B was 90% (v/v) acetonitrile containing 0.1% (v/v) formic acid. Peptides were eluted at a flow rate of 300 nl/min with a linear gradient from 2.2% to 14% B over 22 min, followed by 14% to 33% B over the next 22 min (22–44 min). The column was then washed by ramping to 100% B within 5 min and holding for 8 min. Data were acquired in positive ion polarity mode using data-independent acquisition coupled with parallel accumulation-serial fragmentation (dia-PASEF). Ion mobility was scanned from 0.70 to 1.30 Vs/cm² (1/K₀) with a ramp time of 100 ms. Each acquisition cycle consisted of one MS1 scan (m/z 100–1700) followed by eight dia-PASEF ramps. Each ramp contained three isolation windows, for a total of 24 variable-width windows. These windows covered m/z 350–1000 and 1/K₀ 0.70–1.30 Vs/cm² without gaps or overlaps. Window placement was optimized with py_diAID [35] using a DIA-NN [36] precursor report of a pooled *Chlamydomonas reinhardtii* digest. The complete window scheme is provided in Supplementary Table S3. Collision energy was ramped linearly with ion mobility from 20 eV at 1/K₀ = 0.6 Vs/cm² to 59 eV at 1/K₀ = 1.6 Vs/cm². High-sensitivity detection was enabled. The mass spectrometry proteomics data have been deposited to the ProteomeXchange Consortium via the PRIDE partner repository [37, 38] with the dataset identifier PXD087595.

#### Data processing and bioinformatics

Raw data were processed with DIA-NN 2.6.1 [36] in library-free mode. A predicted spectral library was generated *in silico* from the human UniProt reference proteome (UP000005640, downloaded 9 July 2026), supplemented with the cRAP contaminant database (camprotR version, 2019-04-01). The *in silico* digest assumed trypsin specificity (cleavage after K/R) with up to one missed cleavage, a peptide length of 7–30 residues, precursor charge states 1–4, a precursor m/z range of 300–1800 and a fragment m/z range of 200–1800. Carbamidomethylation of cysteine was set as a fixed modification, and N-terminal methionine excision was enabled. No variable modifications were considered. MS1 and MS2 mass accuracies were fixed at 15 ppm. Match-between-runs was enabled: in a first pass, an empirical spectral library was generated from the DIA data using the predicted library, and all runs were then reanalyzed with this experiment-specific library in a second pass. Contaminant peptides were excluded from normalization and quantification. All other parameters were left at their defaults. Precursors and protein groups were filtered at 1% false discovery rate (FDR).

Differential abundance analysis was carried out on the DIA-NN protein group output using DIA-Analyst (v0.11, https://analyst-suites.org/apps/dia-analyst/). Following contaminant filtering, protein intensity values were subjected to variance stabilizing normalization (vsn), and missing values were imputed using a minimal probability (MinProb) approach, i.e., by random draws from a Gaussian distribution centered on a low quantile of the observed intensities. Differential abundance was assessed with limma-based [39] moderated t-tests, and *p*-values were corrected for multiple testing using the Benjamini–Hochberg procedure. Proteins with an adjusted *p*-value<0.01 and an absolute log_2_ fold change≥2 were considered significantly differentially abundant.

Functional biological interpretation, including Gene Ontology (GO) enrichment and Gene Set Enrichment Analysis (GSEA), was performed using WebGestalt [40]. For GSEA, the entire quantified proteome was ranked by log_2_ fold change and mapped against the Reactome pathway database to determine Normalized Enrichment Scores (NES) and extract leading-edge proteins.

### QUANTIFICATION AND STATISTICAL ANALYSIS

All quantitative data are presented as mean ± standard deviation (SD) of at least three independent experiments unless otherwise indicated. Statistical analyses and curve fittings were performed using OriginPro (v2026b, OriginLab). Normality was assessed using the Shapiro-Wilk test. Group comparisons were evaluated using one-way analysis of variance (ANOVA) with Scheffé’s *post-hoc* test for normally distributed datasets, or the Kruskal-Wallis test followed by Dunn’s *post-hoc* test for non-parametric data. Statistical significance was defined as p<0.05 (\**p*≤0.05, \*\**p*≤0.01, \*\*\**p*≤0.001).

