## Supplementary material for "F_1_F_O_-ATP synthase subunit DAPIT rewires mitochondrial metabolism in cancer cells to drive cellular hyperproliferation": Manuscript: Supplemental Information.pdf

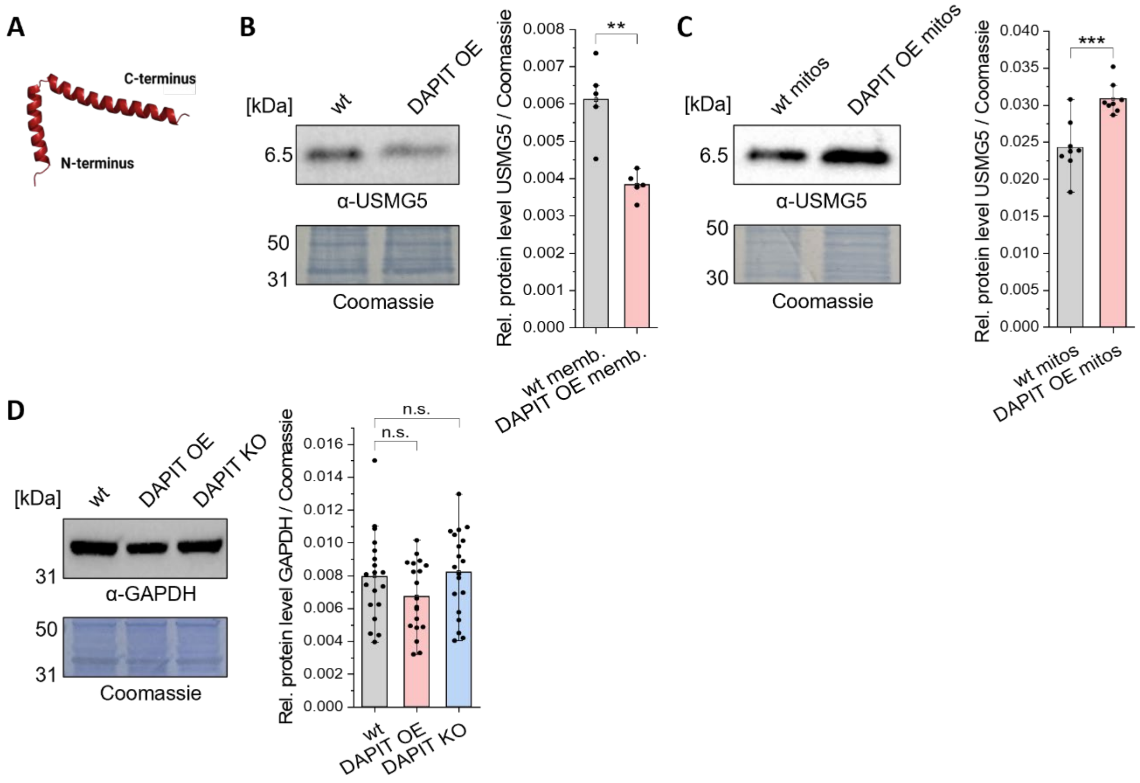

**Figure S1: Subcellular localization of overexpressed DAPIT and validation of GAPDH as reference housekeeping protein.**
**(A)** Predicted human DAPIT protein structure using AlphaFold 3 [21].
**(B)** Representative Western blot and quantification of relative (rel.) protein levels of USMG5 (also known as ATP synthase subunit DAPIT) in membrane fractions of HeLa wildtype (wt), DAPIT overexpressing (OE), and DAPIT knockout (KO) cells normalized to Coomassie staining (N=3, n=2; one-way ANOVA with Scheffé's *post-hoc* test).
**(C)** Representative Western blot and quantification of relative protein levels of USMG5 (DAPIT) in isolated mitochondria of HeLa wt, DAPIT OE, and DAPIT KO normalized to Coomassie staining (N=4, n=2; one-way ANOVA with Scheffé's *post-hoc* test; independent mitochondrial isolations).
**(D)** Representative Western blot and quantification of relative protein levels of GAPDH, normalized to Coomassie staining, showing stable GAPDH expression across all cell lines and confirming its suitability as loading control (N=6, n=3; one-way ANOVA with Scheffé's *post-hoc* test).
Data are presented as mean ± SD of N=3-6 independent biological experiments with overlaid individual data points. The exact number of independent experiments (N) and each number of analyzed technical replicates, wells, or individual cells (n) are indicated in the respective brackets for each panel Statistics: ***$p \leq 0.001$; **$p \leq 0.01$; *$p \leq 0.05$. ANOVA, analysis of variance; DAPIT, diabetes-associated protein in insulin-sensitive tissue; GAPDH, glyceraldehyde-3-phosphate dehydrogenase; SD, standard deviation; USMG5, up-regulated during skeletal muscle growth protein 5.

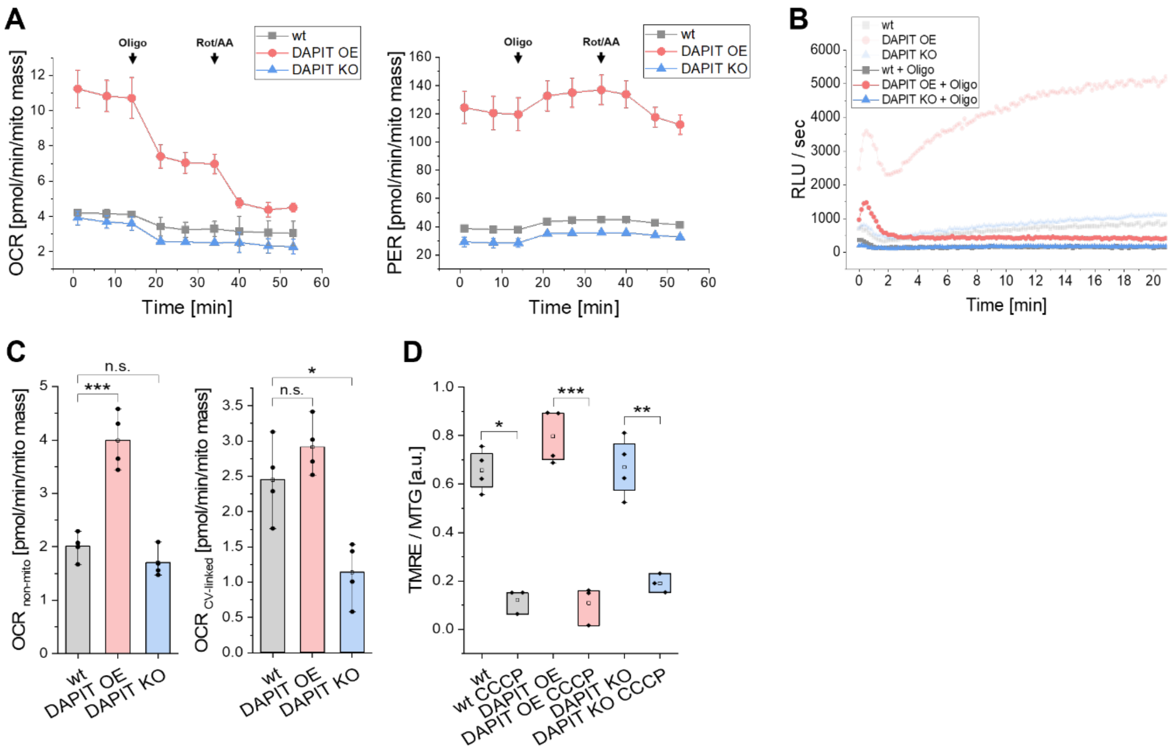

**Figure S2: Bioenergetic profiling and mitochondrial functional analysis of DAPIT overexpressing and knockout cells.**
**(A)** Oxygen consumption rates (OCR) and proton efflux rates (PER) of HeLa wildtype (wt), DAPIT overexpressing (OE), and DAPIT knockout (KO) cells were determined with an automatic Seahorse XF96 flux analyzer and normalized to mitochondrial mass (mito mass). ATP production rates were determined by subsequently adding ATP synthase inhibitor oligomycin (Oligo, 1 µM), complex I inhibitor rotenone (Rot, 1 µM), and complex III inhibitor antimycin A (AA, 1 µM), as indicated by arrows (N=3, n=20 wells).
**(B)** Representative kinetic profile of ATP production rates in freshly isolated mitochondria, measured via a luciferase assay in the absence/presence of ATP synthase inhibitor oligomycin (Oligo, 10 µM) to validate the mitochondrial contribution to ATP synthesis. Equal amounts of total protein (50 µg mitochondria) were used. ATP production was fueled via complex II by addition of succinate (5 mM) in presence of rotenone (5 mM) to inhibit complex I. ADP (4 mM) was added as a further substrate and adenylate kinase was inhibited with Ap5A (20 µM) (N=3, n=2 wells; independent mitochondrial isolations).
**(C)** OCR of HeLa wt, DAPIT OE, and DAPIT KO cells were determined with an automatic Seahorse XF96 flux analyzer and normalized to mitochondrial mass. Non-mitochondrial (non-mito) and ATP synthase (CV)-linked respiration were determined (N=4, n=20 wells; one-way ANOVA with Scheffé's *post-hoc* test).
**(D)** Quantification of mitochondrial membrane potential ($\Delta\Psi_m$) using live-cell TMRE staining, calculated as the TMRE/MTG ratio to normalize for mitochondrial mass (determined via MitoTracker Green, MTG). As control, uncoupler CCCP (10 µM) was added after staining (N=4, n=35 single cells; CCCP N=3, n=21 single cells; wt Kruskal-Wallis with Dunn's *post-hoc* test; DAPIT OE and KO one-way ANOVA with Scheffé's *post-hoc* test).
Data are presented as mean ± SD of N=3-4 independent biological experiments. The exact number of independent experiments (N) and each number of analyzed technical replicates, wells, or individual cells (n) are indicated in the respective brackets for each panel. Statistics: ***p≤0.001; **p≤0.01; *p≤0.05. ANOVA, analysis of variance; Ap5A, diadenosine pentaphosphate; CCCP, carbonyl cyanide *m*-chlorophenyl hydrazone; DAPIT, diabetes-associated protein in insulin-sensitive tissue; RLU, relative light units; SD, standard deviation; TMRE, tetramethylrhodamine ethyl ester.

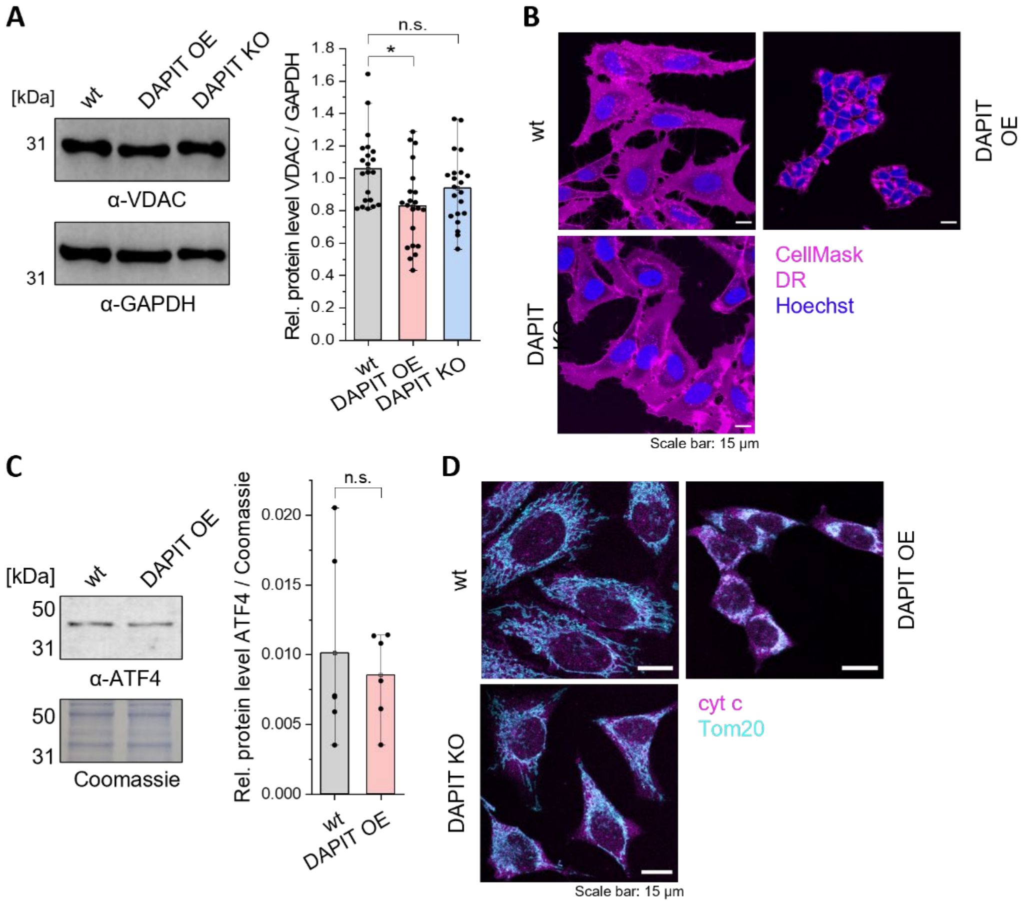

**Figure S3: Effects of DAPIT overexpression on cellular/nuclear morphology, ATF4-mediated stress signaling, and apoptotic cytochrome c release.**

**(A)** Representative Western blot and quantification of relative (rel.) protein levels of VDAC, used as marker for mitochondrial mass. Protein levels were normalized to GAPDH (N=5, n=4; Kruskal-Wallis with Dunn's *post-hoc* test).

**(B)** Representative overlaid images from live-cell fluorescence microscopy showing plasma membrane staining with CellMask Deep-Red (DR, magenta) and cell nuclei with Hoechst 33342 (blue) to quantify alterations in cell area and nuclear size.

**(C)** Representative Western blot and quantification of relative protein levels of ATF4 in HeLa wildtype (wt), DAPIT overexpressing (OE), and DAPIT knockout (KO) cells normalized to Coomassie staining (N=3, n=2; one-way ANOVA with Scheffé's *post-hoc* test).

**(D)** Representative overlaid images from immunofluorescence staining of cytochrome c (cyt c, magenta) and mitochondrial staining with TOM20 (cyan). Loss of cytochrome c/TOM20 colocalization serves as an indicator of mitochondrial outer membrane permeabilization and apoptotic cytochrome c release.

Data are presented as mean ± SD of N=3-5 independent biological experiments with overlaid individual data points. The exact number of independent experiments (N) and each number of analyzed technical replicates, wells, or individual cells (n) are indicated in the respective brackets for each panel. Statistics: ***p≤0.001; **p≤0.01; *p≤0.05. ANOVA, analysis of variance; ATF4, activating transcription factor 4; DAPIT, diabetes-associated protein in insulin-sensitive tissue; GAPDH, glyceraldehyde-3-phosphate dehydrogenase; SD, standard deviation; TOM20, translocase of the outer mitochondrial membrane 20; VDAC, voltage-dependent anion channel.

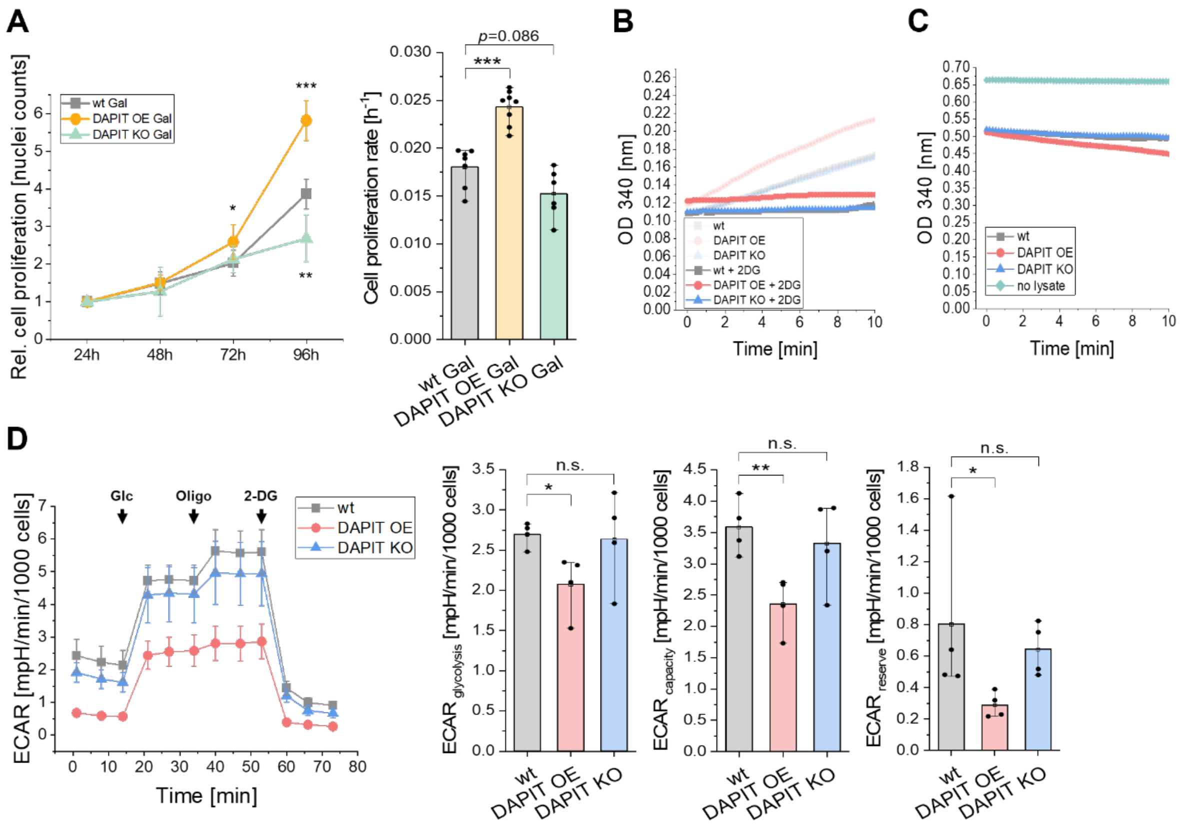

**Figure S4: Cell proliferation under OXPHOS conditions, glycolytic profiling, and functional validation of coupled enzyme assays.**

**(A)** Growth curves of HeLa wildtype (wt), DAPIT overexpressing (OE), and DAPIT knockout (KO) cells under OXPHOS conditions (galactose medium, Gal) over a period of 96 h. Cell numbers were determined every 24 h by counting Hoechst 33342-stained nuclei using a fluorescence microplate reader. To account for initial seeding differences, cell numbers were normalized to 24 h timepoint. Cell proliferation rates were calculated from the slope of exponential growth phase (N=3, n=3; one-way ANOVA with Scheffé's *post-hoc* test).

**(B)** Representative kinetic profile of hexokinase 2 (HK2) activity from fresh cell lysates in the absence/presence of 2-deoxyglucose (2-DG, 150 mM), monitored by the accumulation of NADH over time. 2-DG was added as competitive inhibitor of HK2 to confirm assay specificity. The assay was coupled to an excess of exogenous glucose-6-phosphate dehydrogenase to continuously convert HK2-generated glucose-6-phosphate. Reduction of $NAD^+$ to NADH was measured in real-time via absorbance (optical density, OD) at 340 nm using equal amounts of total protein (50 µg cell lysate) (N=3, n=3 wells).

**(C)** Representative kinetic profile of endogenous pyruvate consumption in the absence/presence of fresh cell lysates, monitored by depletion of NADH. Background control for coupled enzyme assay was performed in absence of cell lysates to determine baseline substrate interference. The assay was coupled to an excess of exogenous lactate dehydrogenase and NADH to continuously convert endogenous pyruvate into lactate. Oxidation of NADH to $NAD^+$ was measured in real-time via decrease in absorbance at 340 nm using equal amounts of total protein (50 µg cell lysate) (N=3, n=4 wells).

**(D)** Functional test of glycolysis with subsequent addition of saturating amounts of glucose (Glc, 10 mM), ATP synthase inhibitor oligomycin (Oligo, 1 µM), and glycolysis inhibitor 2-deoxy-D-glucose (2-DG, 50 mM). Extracellular acidification rates (ECAR) of HeLa wt, DAPIT OE, and DAPIT KO cells were determined with an automatic Seahorse XF96 flux analyzer and normalized to cell number (1000 cells). Cell number was determined by counting Hoechst 33342-stained nuclei using a fluorescence microplate reader. Glycolysis, glycolytic capacity, and glycolytic reserve were determined (N=4, n=20 wells; glycolysis and capacity one-way ANOVA with Scheffé's *post-hoc* test; reserve Kruskal-Wallis with Dunn's *post-hoc* test).

Data are presented as mean ± SD of N=3-4 independent biological experiments. The exact number of independent experiments (N) and each number of analyzed technical replicates, wells, or individual cells (n) are indicated in the respective brackets for each panel. Individual data points are overlaid in (A). Statistics: ***p≤0.001; **p≤0.01; *p≤0.05. ANOVA, analysis of variance; DAPIT, diabetes-associated protein in insulin-sensitive tissue; $NAD^+$, oxidized nicotinamide adenine dinucleotide; NADH, reduced nicotinamide adenine dinucleotide; SD, standard deviation

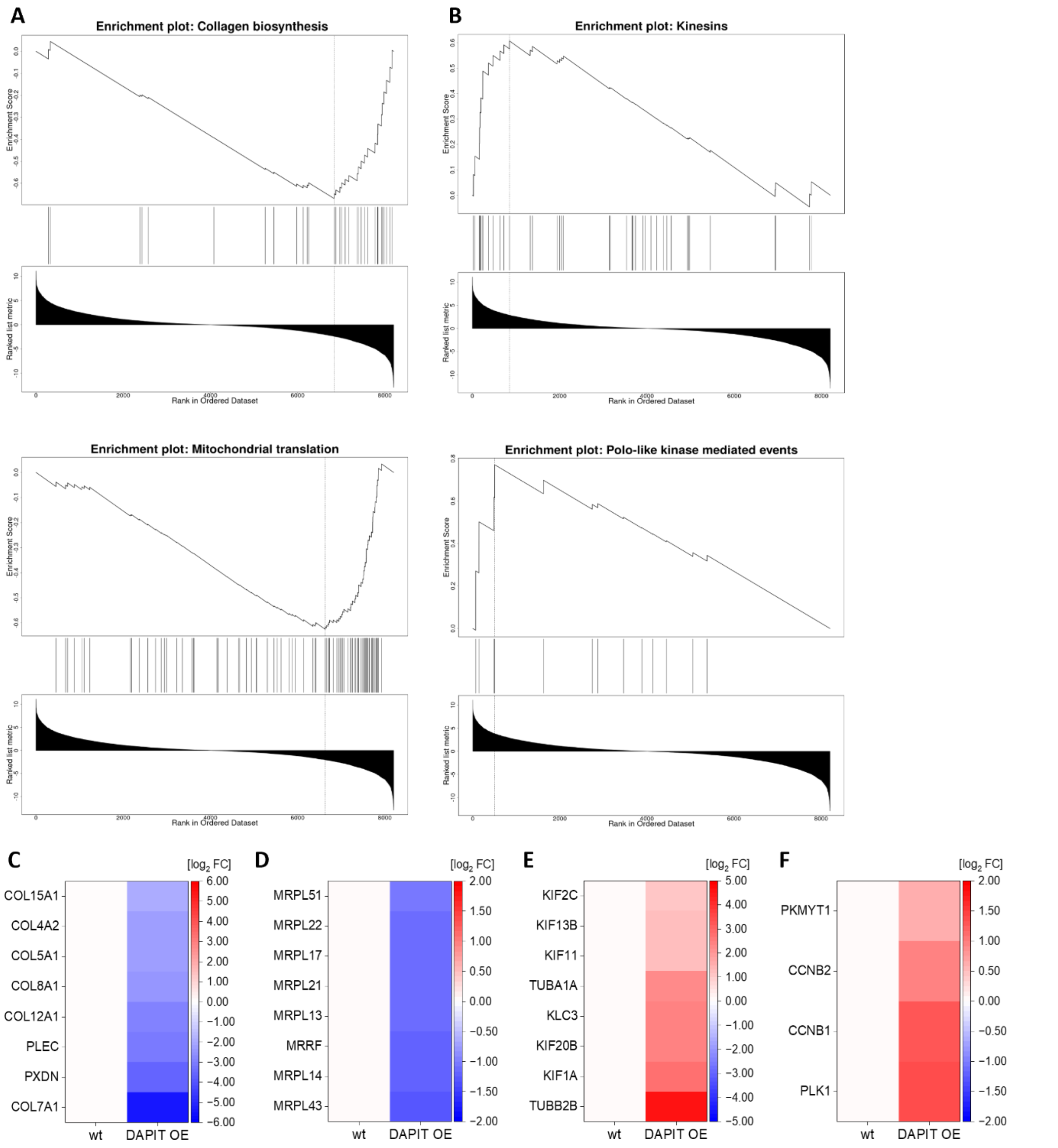

**Figure S5: Detailed Gene Set Enrichment Analysis and leading-edge analysis of DAPIT overexpressing proteome.**
**(A, B)** Gene Set Enrichment Analysis plots for selected **(A)** top downregulated (e.g., collagen biosynthesis and mitochondrial translation) and **(B)** top upregulated (e.g., kinesins and Polo-like kinase mediated events) pathways in DAPIT overexpressing (OE) cells versus HeLa wildtype (wt) cells. Curve (top section) indicates running enrichment score, vertical black lines (middle section) represent position of individual pathway proteins within ranked dataset, and graph (bottom section) shows ranked list metric.
**(C-F)** Heatmaps illustrating $\log_2$ FC of corresponding leading-edge proteins for pathways shown in **(A)** and **(B)** – **(C)** collagen biosynthesis pathway, **(D)** mitochondrial translation pathway, **(E)** kinesins, and **(F)** Polo-like kinase mediated events. Color scale is centered at zero (white, representing wt baseline), with blue indicating downregulation (log2 FC<0) and red indicating upregulation (log2 FC>0) in DAPIT OE cells.
Data were obtained from N=4 independent biological experiments. DAPIT, diabetes-associated protein in insulin-sensitive tissue; FC, fold change.

**Supplementary Table 1:** Primers (fw, forward; rev, reverse), along with their respective sequences.

| **Primers (targets)** | **Sequence** |
|---|---|
| *AMPK* fw | ACA GCC GAG AAG CAG AAA CAC |
| *AMPK* rev | GCC CAG TCA ATT CAT GTT TGC C |
| *Bax* fw | TGG CAG CTG ACA TGT TTT CTG AC |
| *Bax* rev | TCA CCC AAC CAC CCT GGT CTT |
| *Bcl2* fw | TCG CCC TGT GGA TGA CTG A |
| *Bcl2* rev | CAG AGA CAG CCA GGA GAA ATC A |
| *Casp9* fw | GTT TGA GGA CCT TCG ACC AGC T |
| *Casp9* rev | CAA CGT ACC AGG AGC CAC TCT T |
| *GPX7* fw | CCA CTT TAA CGT GCT CGC CTT C |
| *GPX7* rev | AAG CCC CTA CCA CCT TTC CAT C |
| *HK2* fw | TTT ACC TTC TCC TTC CCC TGC C |
| *HK2* rev | TGT CGT TCA CCA CAG CAA CC |
| *mTOR* fw | AGC ATC GGA TGC TTA GGA GTG G |
| *mTOR* rev | CAG CCA GTC ATC TTT GGA GAC C |
| *p16* fw | GAA GGT CCC TCA GAC ATC CCC |
| *p16* rev | CCC TGT AGG ACC TTC GGT GAC |
| *p21* fw | CCG AAG TCA GTT CCT TGT GG |
| *p21* rev | CAT GGG TTC TGA CGG ACA T |
| *p53* fw | CCC CTC CTG GCC CCT GTC ATC TTC |
| *p53* rev | GCA GCG CCT CAC AAC CTC CGT CA[illegible] |
| *PGC1α* fw | CCA AAG GAT GCG CTC TCG TTC A |
| *PGC1α* rev | CGG TGT CTG TAG TGG CTT GAC T |
| *RPLP0* fw | TGG TCA TCC AGC AGG TGT TCG A |
| *RPLP0* rev | ACA GAC ACT GGC AAC ATT GCG G |
| *UCP2* fw | AAG ACC ATT GCC CGA GAG GAA G |
| *UCP2* rev | CAG AAG TGA AGT GGC AAG GGA G |
| *UCP3* fw | GGA GCG ACA GAA AAT ACA GCG G |
| *UCP3* rev | AAA GGC AGA GAC AAA GTG GCA G |

**Supplementary Table 2:** List of primary and secondary antibodies, including host species, manufacturers, and working dilutions.

| Primary Antibody | Host Organism | Manufactures | Dilution |
|---|---|---|---|
| Primary Antibodies for IF | | | |
| anti-cytochrome c | ms | Proteintech (66264-1-Ig) | 1:200 |
| anti-TOM20 | rb | Proteintech (11802-1-AP) | 1:400 |
| Primary Antibodies for SDS- or native-PAGE | | | |
| α-ATP5A | ms | abcam (ab14748) | 1:1000 |
| α-ATP5B | rb | abcam (ab170947) | 1:1000 |
| α-ATP5I | rb | abcam (ab122241) | 1:1000 |
| α-ATPIF1 | rb | Cell Signaling (D6P1Q) | 1:1000 |
| α-ATP5MC1 | rb | abcam (ab181243) | 1:1000 |
| α-CREB2 H-290 (ATP4) | rb | Santa Cruz Biotechnology (sc-22800 | 1:500 |
| α-GAPDH | ms | Proteintech (60004-1-Ig) | 1:10000 |
| α-USMG5 | rb | Proteintech (17716-1-AP) | 1:500 |
| α-VDAC 1/2 | rb | Proteintech (10866-1-AP) | 1:1000 |
| **Secondary Antibody** | **Manufactures** | **Dilution** | |
| Secondary Antibodies for IF | | | |
| Alexa Fluor 647 goat anti-mouse IgG H&L | abcam (ab150115) | 1:600 | |
| Alexa Fluor 488 goat anti-rabbit IgG H&L | abcam (ab150077) | 1:600 | |
| Secondary Antibodies for SDS- or BN-PAGE | | | |
| Goat Anti-Mouse IgG H&L | Biozol (LIN-ITC207P) | 1:2000 | |
| Goat Anti-Rabbit IgG H&L | Biozol (LIN-ITC208P) | 1:2000 | |

**Supplementary Table 3:** dia-PASEF isolation window scheme

| #MS Type | Cycle Id | Start IM [1/K0] | End IM [1/K0] | Start Mass [m/z] | End Mass [m/z] | CE [eV] | Isolation width [m/z] |
|---|---|---|---|---|---|---|---|
| MS1 | 0 | - | - | - | - | - | - |
| PASEF | 1 | 0.7000 | 0.8100 | 350.18 | 395.86 | - | 45.68 |
| PASEF | 1 | 0.8100 | 0.9700 | 549.61 | 568.84 | - | 19.23 |
| PASEF | 1 | 0.9700 | 1.3000 | 722.99 | 748.40 | - | 25.41 |
| PASEF | 2 | 0.7000 | 0.8400 | 395.86 | 422.71 | - | 26.85 |
| PASEF | 2 | 0.8400 | 0.9900 | 568.84 | 589.35 | - | 20.51 |
| PASEF | 2 | 0.9900 | 1.3000 | 748.40 | 775.92 | - | 27.52 |
| PASEF | 3 | 0.7000 | 0.8600 | 422.71 | 445.75 | - | 23.04 |
| PASEF | 3 | 0.8600 | 1.0000 | 589.35 | 609.33 | - | 19.98 |
| PASEF | 3 | 1.0000 | 1.3000 | 775.92 | 805.71 | - | 29.79 |
| PASEF | 4 | 0.7000 | 0.8800 | 445.75 | 467.25 | - | 21.50 |
| PASEF | 4 | 0.8800 | 1.0200 | 609.33 | 630.34 | - | 21.01 |
| PASEF | 4 | 1.0200 | 1.3000 | 805.71 | 836.95 | - | 31.24 |
| PASEF | 5 | 0.7000 | 0.8900 | 467.25 | 489.22 | - | 21.97 |
| PASEF | 5 | 0.8900 | 1.0300 | 630.34 | 651.66 | - | 21.32 |
| PASEF | 5 | 1.0300 | 1.3000 | 836.95 | 870.48 | - | 33.53 |
| PASEF | 6 | 0.7000 | 0.9100 | 489.22 | 509.77 | - | 20.55 |
| PASEF | 6 | 0.9100 | 1.0500 | 651.66 | 674.34 | - | 22.68 |
| PASEF | 6 | 1.0500 | 1.3000 | 870.48 | 908.48 | - | 38.00 |
| PASEF | 7 | 0.7000 | 0.9300 | 509.77 | 529.75 | - | 19.98 |
| PASEF | 7 | 0.9300 | 1.0600 | 674.34 | 698.90 | - | 24.56 |
| PASEF | 7 | 1.0600 | 1.3000 | 908.48 | 952.94 | - | 44.46 |
| PASEF | 8 | 0.7000 | 0.9500 | 529.75 | 549.61 | - | 19.86 |
| PASEF | 8 | 0.9500 | 1.0800 | 698.90 | 722.99 | - | 24.09 |
| PASEF | 8 | 1.0800 | 1.3000 | 952.94 | 999.98 | - | 47.04 |
